# Supergene evolution via gain of autoregulation

**DOI:** 10.1101/2024.01.09.574839

**Authors:** Nicholas W. VanKuren, Sofia I. Sheikh, Claire L. Fu, Darli Massardo, Wei Lu, Marcus R. Kronforst

## Abstract

Development requires the coordinated action of many genes across space and time, yet numerous species have evolved the ability to develop multiple discrete, alternate phenotypes^1–5^. Such polymorphisms are often controlled by supergenes, sets of tightly-linked loci that function together to control development of a polymorphic phenotype^6–10^. Although theories of supergene evolution are well-established, the mutations that cause functional differences between supergene alleles have been difficult to identify. The *doublesex* gene is a master regulator of insect sexual differentiation but has been co-opted to function as a supergene in multiple *Papilio* swallowtail butterflies, where divergent *dsx* alleles control development of discrete non-mimetic or mimetic female wing shapes and color patterns^11–15^. Here we demonstrate that the *Papilio alphenor* supergene evolved via recruitment of six new *cis*-regulatory elements (CREs) that control allele-specific *dsx* expression. Most *dsx* CREs, including four of the six new CREs, are bound by the DSX transcription factor itself. Our findings provide experimental support to classic supergene theory and suggest that autoregulation may provide a simple route to supergene origination and to the co-option of pleiotropic genes into new developmental roles.

---

Supergenes are predicted to evolve through a series of four stages^16,17^. First, a mutation arises that causes development of a novel, weakly adaptive phenotype. A second mutation then arises that improves the adaptive phenotype. Selection for the improved phenotype favors the evolution of linkage disequilibrium between the mutations and reduced recombination between the new supergene alleles. Linkage disequilibrium, caused by a variety of mechanisms including inversions, allows each allele to accumulate additional mutations that refine the supergene’s function and therefore the novel adaptive phenotype^6,7,10,16^. In addition, unlinked epistatic modifier loci may evolve to further refine the adaptive phenotype^8^.

While sex chromosomes are perhaps the most well-known supergenes, complex balanced polymorphisms including alternative social structures in ants^2,18^, flower structures^7^, and bird mating morphs^3^ have now been traced back to allelic variation in supergenes. Recent genomic studies show that supergenes often span multiple megabases, contain tens to hundreds of genes, and exhibit significantly reduced recombination^19^. Yet these same features have made it difficult to identify the genetic variants that cause functional differences between alleles, and therefore the evolutionary origins of the alleles, their regulatory architectures, and the polymorphisms they control.

Here we aimed to identify the functional genetic basis of supergene mimicry in *Papilio* swallowtail butterflies^6,20–22^. *doublesex* functions as a supergene in at least five species of *Papilio* swallowtail butterflies, where it controls development of discrete female wing color patterns^11–14^. The switch from a male-like non-mimetic color pattern to a novel mimetic color pattern in the closely related species *P. alphenor* and *P. polytes* is caused by the novel *H* allele^11,12,14^. The *H* allele is inverted relative to the ancestral *h* allele, has an extremely divergent sequence from *h*, and contains *dsx* and a novel non-coding gene, *untranslated three exons* (*U3X*; **Fig 1A-B**)^11–13^. A unique spike of *dsx^H^* expression in early pupal wings initiates the mimetic color development program, but the genetic basis of this novel expression pattern remained unknown (**Fig 1C**)^15^.

**Fig 1.**
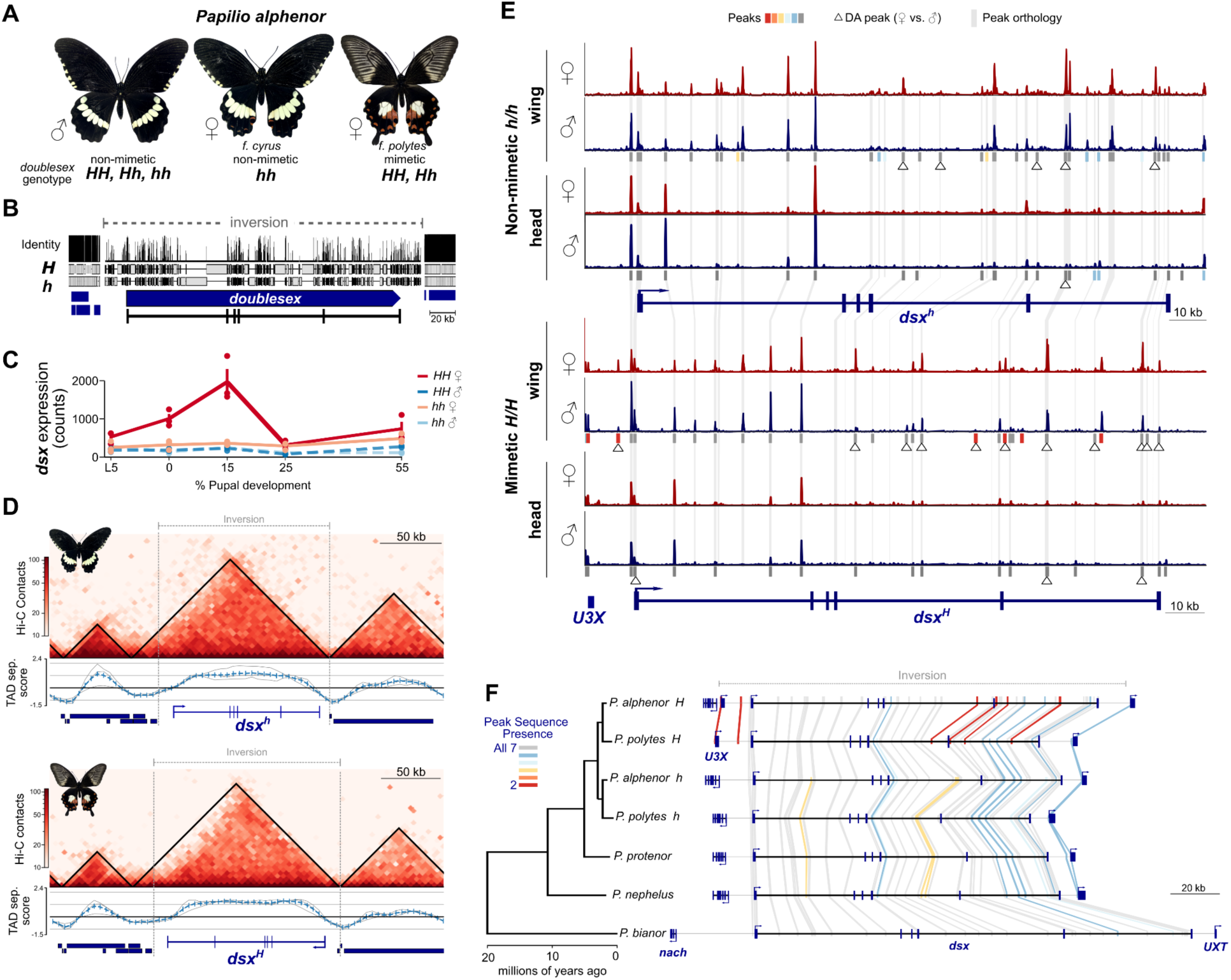
Regulatory architecture of the *doublesex* supergene. (**A**) Female-limited mimicry polymorphism in *Papilio alphenor*. (**B**) Pairwise alignment of *dsx* supergene alleles and flanking 20 kb. The mimetic *H* allele is un-inverted for alignment and display. (**C**) *dsx* hindwing expression in each group across development^15^. (**D**) Hi-C contacts, topologically associating domains (TADs), and TAD separation scores near the inversion in early pupal female wings. Gene models are shown along the x axis. (**E**) Transposase-accessible chromatin within the inversion in early pupal (15% pupal development) wings and heads. All tracks are normalized and shown on the same scale [0-450]. Peak calls and differentially accessible (DA) peaks between females and males are shown below. Orthologous peak sequences between the alleles are connected by shaded lines. Additional information and replicates are shown in Supplementary Figures 2-5 and Supplementary Table 1. Peak color coding corresponds to **F**. (**F**) Orthology and synteny between *P. alphenor* ATAC-seq peaks and outgroup alleles. Orthology was determined by BLASTing *P. alphenor* peak sequences to the genome regions bounded by *nach* and *UXT,* which flank the inverted region, in each species. The *P. alphenor* and *P. polytes H* alleles are un-inverted for display. Peak conservation (i.e. the number of sequences in which the peak sequence is present) is denoted by color and lines connect orthologous regions in each allele. Lines are thickened to make them easier to view. Additional details and results can be found in the Methods and Supplementary Tables 2 - 7.

## *The mimetic* H *allele gained multiple novel CREs*

We therefore began our investigations by characterizing the *cis*-regulatory architecture of the *P. alphenor* supergene in the developing wing (**Fig 1**). We first narrowed our search for CREs controlling *dsx* wing expression using Hi-C to identify topologically associating domains (TADs) containing each allele, which we expected to define the local regulatory region^23^. Each allele was contained within a single TAD harboring *dsx* and the adjacent genes *sir-2, rad51*, and *nach*; the mimetic *H* TAD additionally contained *U3X* (**Fig 1D**). While the right TAD boundaries coincided with the right inversion breakpoint in both alleles, the left TAD boundaries were 21.9 kb and 17.9 kb outside of the inversion, suggesting that the inversion itself caused minor changes to the local topology despite high sequence and structural divergence between the alleles (**Fig 1B**). TAD boundaries were similar in early pupal male wings and mid-pupal male and female wings (Supplementary Figure 1). We therefore expected CREs controlling *dsx* wing expression to be within or just to the left of the inversion.

We identified potential *dsx* CREs using the assay for transposase-accessible chromatin (ATAC) on early and mid-pupal wings and heads from homozygous females and males (**Fig 1E**; Supplementary Figures 2-5; Supplementary Tables 1-3)^15,24^. We first focused on early pupal development, where *dsx^H^* expression spikes in mimetic female wings (**Fig 1C**)^15^. The mimetic *H* allele contained 28 ATAC peaks in early pupal wings, with most peaks within or near *dsx^H^*: 22 within introns plus peaks within exon 1, exon 6, and 0.47 kb and 3.9 kb upstream of the *dsx^H^*promoter. Peaks were also found at the *UXT* and *U3X* promoters. The non-mimetic *h* allele contained 33 peaks in early pupal wings. These *h* peaks were located in similar relative locations to *H* peaks, but were absent from ∼4 kb upstream of *dsx^h^* (**Fig 1E**). We considered peaks outside of the *UXT, U3X,* and *dsx* promoters as potential CREs.

The different numbers of peaks in the two alleles suggested that allele-specific CREs could contribute to their functional differences, but the extreme sequence divergence between the alleles precluded direct comparisons between peaks. We therefore identified orthologous *h* and *H* peaks and polarized peak gain and loss using BLAST^25^ to search for peak sequences in *P. alphenor, P. polytes* and three additional, monomorphic *Papilio* species (**Fig 1F**; Supplementary Tables 4-5). Despite the enormous sequence divergence between the *h* and *H* alleles (**Fig 1B**), we found high conservation of CRE sequences and synteny over 20 million years of evolution. Importantly, all *h* peak sequences were found in the *P. polytes h* allele and at least two other species. However, six *H* peaks -- the *U3X* promoter and five CREs -- were unique to the *H* allele and shared by both *P. alphenor* and *P. polytes*, strongly suggesting that the *H* allele gained multiple novel *dsx* CREs (**Fig 1F**). Both alleles control sexual differentiation equally well, so these *H*-specific CREs are unlikely to be essential for *dsx^H^* expression in other tissues. All *H*-specific CREs were active in early pupal wings but absent in head and mid-pupal wings, suggesting that these CREs are specifically involved in the unique spike of *dsx^H^*expression in early pupal mimetic female wings (**Fig 1C**; Supplementary Figure 4).

This hypothesis was supported by patterns of differential CRE accessibility across the supergene (**Fig 1F**; Supplementary Tables 6-7). We found that 44% (11/25) of *H* CREs were differentially accessible (DA) between the sexes, including 60% (3/5) of *H*-specific CREs. In contrast, only 17% (5/29) of *h* CREs were DA. The smaller number of DA peaks in the *h* allele is consistent with the similar *dsx* expression patterns and small amount of dimorphism between non-mimetic females and males (**Fig 1E**)^26^. This sexually dimorphic accessibility is likely involved in dimorphic color pattern development, as no CREs were DA specifically in heads and only one was DA in both heads and wings.

*dsx^H^* thus appeared to be regulated through a small number of *H-*specific CREs. We tested this idea by knocking out *H-*specific CREs (**Fig 2**; Supplementary Figure 6; Supplementary Tables 8 - 9). We expected that knocking out CREs required for *dsx^H^* expression would cause mimetic females to develop non-mimetic patterns. We injected Cas9 and two to four single guide RNAs (sgRNAs) targeting a single CRE into heterozygous eggs, allowed them to develop, and screened adults for mosaic color patterns^27^. Consistent with their role in the mimicry switch, injections targeting 4/5 *H*-specific CREs yielded mimetic females with mosaic patches of non-mimetic pattern (**Fig 2**; Supplementary Figure 6). Mosaic knockouts (mKOs) of each CRE affected all mimetic color pattern elements, including: scale color in light patches, the size and color of submarginal hindwing spots, and the presence of stripes and marginal spots on the forewing (**Fig 2**). These results stand in contrast to knockouts of CREs for genes like *WntA* or *Optix* that pre-figure certain color patterns throughout wing development, which often result in modular changes to specific color pattern elements^27,28^. This difference is likely due to the fact that the *dsx* CREs we tested are controlling the wing-wide spike of *dsx^H^* expression in early pupal wings rather than region-specific expression. In addition, long-read whole genome sequencing of three mKO butterflies strongly suggested that mKO phenotypes were specifically caused by knockout of the target CRE, rather than disruption of the entire gene or multiple CREs (**Fig 2G**; Supplementary Text; Supplementary Figure 7). We conclude that the *dsx* supergene evolved through recruitment of multiple novel CREs spread across 150 kb that enhanced *dsx* expression in early pupal wings to trigger development of a novel mimetic wing pattern.

**Fig 2.**
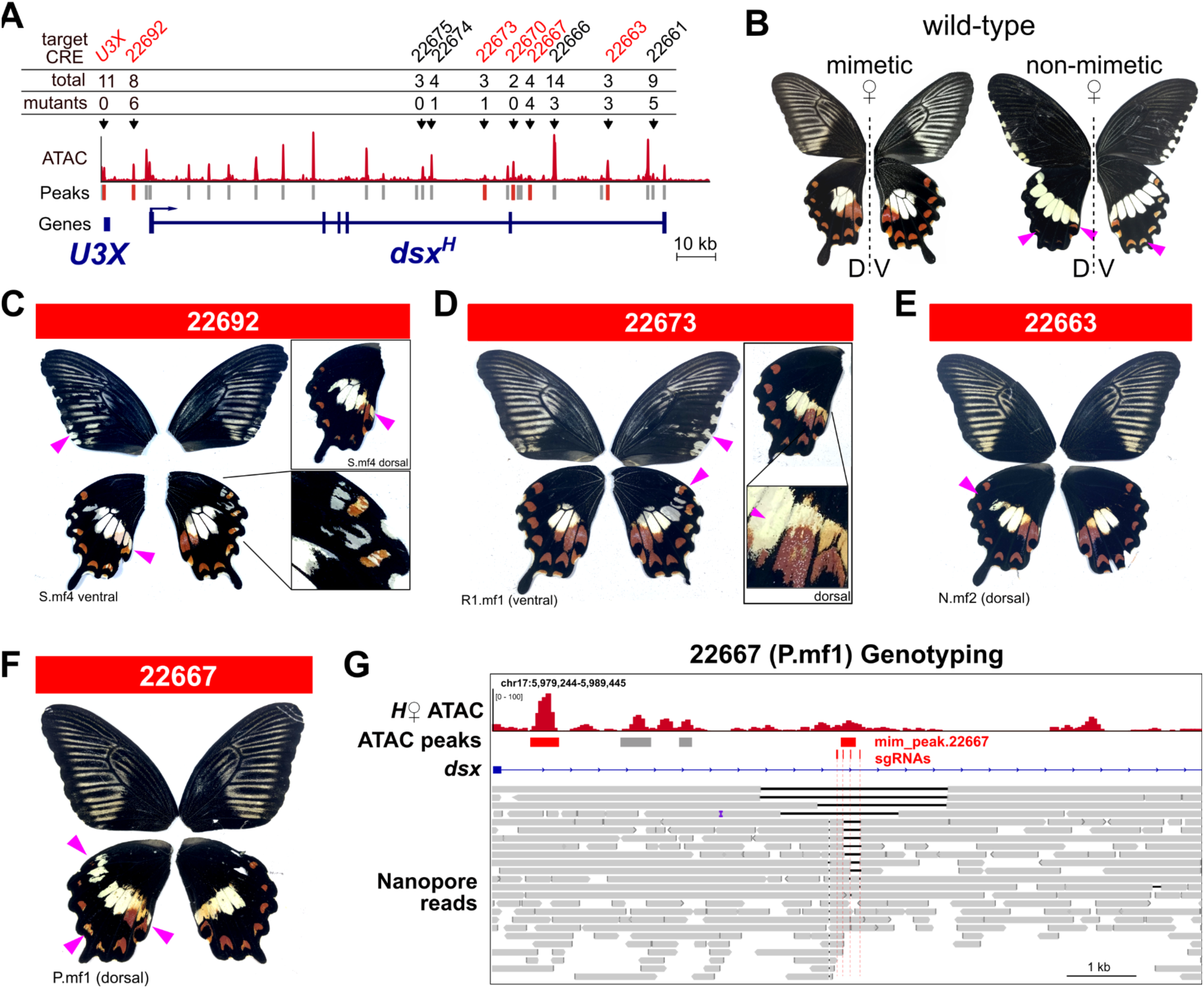
CRISPR/Cas9 knockouts show multiple *H*-specific CREs are required for mimetic color pattern development. **A,** Candidate CREs were targeted for deletion by injecting 2 - 4 sgRNAs and Cas9 into fresh, heterozygous *Hh* eggs (Supplementary Tables 8-9). We targeted all six *H*-specific CREs and four conserved CREs individually. The total numbers of mimetic females (total) and mosaic knockout (mKO) females are shown for each target CRE. We never observed mutant phenotypes in males resulting from these injections. CRE orthology and differential accessibility in early pupal wings are from Fig 1. **B,** Wild-type female color patterns, showing dorsal (D) and ventral (V). Mosaic KO females were identified by the appearance of non-mimetic color patterns in otherwise mimetic females. **C-F,** Examples of mKO females recovered from injections targeting CREs 22692 (**C**), 22673 (**D**), and 22663 (**E**), or 22667 **(F)**. Sample identifiers and surfaces are shown below images. mKOs of additional, conserved CREs can be found in Supplementary Figure 6. **G,** Whole-genome long-read sequencing from the individual shown in **(F)**, demonstrating the range of deletions recovered in a single individual. Of note is the absence of deletions affecting CDS or CREs beyond the target CRE 22667. Indels and variation < 5 bp long are not highlighted. Black lines within read alignments indicate deletions relative to the reference. ATAC data from early pupal mimetic (*H*) female wing is shown, with peak calls below. *Dsx* exon 5 and intron 5 are shown.

## doublesex *is autoregulated*

What transcription factor(s) (TFs) control *dsx* wing expression? The TFs that directly control *dsx* transcription are unknown in any organism. Genetic analyses in *Drosophila* showed that Hox and other TFs, including Sex-combs reduced, Abdominal-B, and Caudal^29^, are required for DSX expression in certain contexts, but it is unknown if these TFs directly regulate *dsx* transcription. On the other hand, ChIP-seq experiments identified binding sites for over 230 different DNA binding proteins within *dsx* in whole adult *Drosophila*, chiefly DSX itself (25 sites)^30^. We attempted to identify TFs that regulate *dsx* expression in the *P. alphenor* wing by searching for known TF binding site motifs enriched in *dsx^H^*CREs using HOMER^31^ (Supplementary Figure 8). CREs were most significantly enriched with motifs for Paired (*p* = 1e- 13), Caudal (*p* =1e-11), Extradenticle (*p* = 1e-10), and DSX (*p* = 1e-9), supporting the idea that *dsx* could be autoregulated.

To test this hypothesis and identify the direct targets of DSX in the wing, we assayed genome-wide patterns of DSX binding in early and mid-pupal wings using CUT&RUN (**Fig 3**; Supplementary Figures 9-11; Supplementary Table 10)^32^. We found 10,318 DSX peaks genome-wide among all samples, with the majority of peaks (82.8%) initially called only in *HH* female samples, where DSX expression is highest. DSX peaks mostly overlapped ATAC peaks (88.2%), and were enriched with a motif similar to the known *Drosophila* DSX binding site (**Fig 3**; *p* = 1e-744; Supplementary Figure 10), supporting the quality of the data. For comparison, 82.2% (9480/11533) H3K4me3 CUT&RUN peaks overlapped ATAC peaks. In addition, we found strong DSX peaks in genes known to be bound by DSX in *Drosophila*, including *bric-a- brac 1* (Supplementary Figure 12).

**Fig 3.**
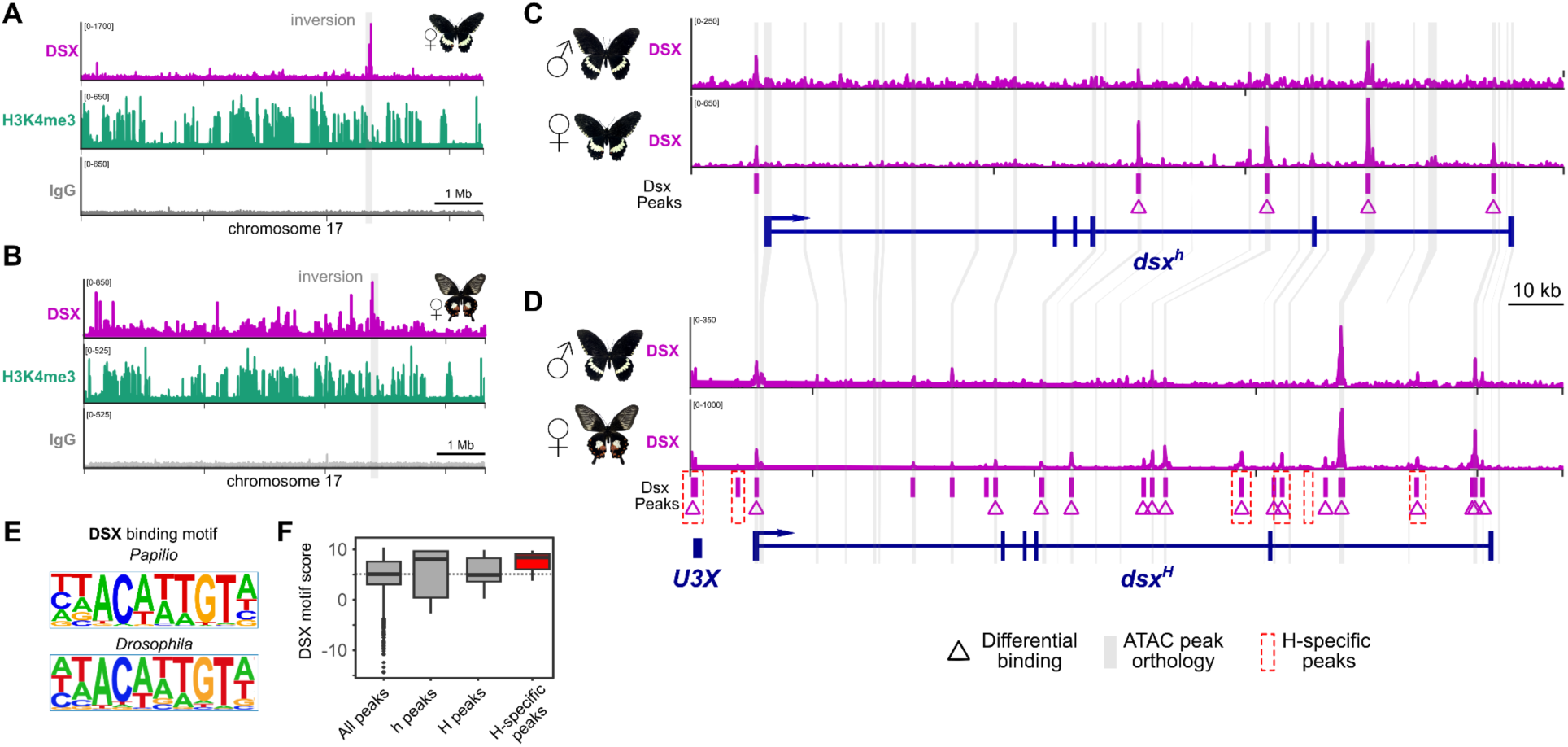
*dsx^H^* gained multiple novel CREs bound by DSX itself. (**A-B**) Normalized coverage tracks for DSX CUT&RUN, the active promoter/enhancer histone mark H3K4 tri-methylation, and negative control immunoglobulin in non-mimetic *hh* (**A**) and mimetic *HH* (**B**) early pupal wings. The whole of chromosome 17 is shown. (**C-D**) DSX binding in the *h* allele (**C**) and *H* allele (**D**) in early pupal wings. DSX peaks, differentially bound (DB) peaks between males and females, and ATAC peak orthology from Fig 1 are shown below. (**E**) The *Papilio* DSX binding motif relative to the *Drosophila* DSX binding motif. Letter heights are proportional to the base frequencies at each position. (**F**) The distribution of log-odds probability scores for DSX motifs in DSX CUT&RUN peaks. Higher values indicate better matches. Scores are shown for motifs in all peaks genome-wide (all peaks); peaks within orthologous *h* and *H* CREs; or in *H-*specific CREs. No pairwise comparisons were significantly different (all Welch’s *t*-test *p*-values >0.1).

Consistent with autoregulation, we found multiple DSX peaks within and just upstream of *dsx* in all samples (**Fig 3**). We found DSX bound to five sites in the non-mimetic *h* allele in early pupal wings, all within conserved CREs. DSX was bound to an additional 20 sites in mimetic *H* allele, including 22 intronic sites, one 0.8 kb and one 4.1 kb upstream of *dsx^H^*, and one at the *U3X* promoter (**Fig 3D**). In addition to the *U3X* promoter, 4/5 *H*-specific CREs were bound by DSX, including three CREs that yielded strong CRISPR/Cas9 mKO phenotypes (**Fig 2**). These results strongly suggest that both alleles are autoregulated, but that the mimetic allele gained multiple novel autoregulatory interactions (**Fig 3D**).

Most DSX binding in the *H* allele seems to be involved in the female-limited polymorphism rather than sexual dimorphism. Four of five *h* DSX peaks were significantly differentially bound (DB) between non-mimetic females and males in early pupal wings (**Fig 3C**).

All four of these peaks were also DB in the *H* allele, suggesting that these CREs mediate sexual dimorphism. An additional 16 DSX peaks were DB between mimetic females and males, including 3/5 *H-*specific CREs, and we predict that these peaks mediate the mimicry switch (**Fig 3C, D**).

The different patterns of DSX binding in conserved CREs appears to be caused by differential use of those CREs rather than mutations that disrupt binding sites, as the binding site sequences are well-conserved between the supergene alleles and across species (**Figs 3E- F**). Log-odds probabilities, measurements of the strength of matches between predicted DSX binding site sequences and the consensus motif, were not significantly different between genome-wide DSX peaks, conserved peaks, or *H-*specific peaks (**Fig 3F**; all Welch’s *t-*test *p-* values > 0.10). In fact, two of the top three matches are found in *H*-specific CREs (Supplementary Table 11). These results suggest that the mimetic *H* allele has gained multiple new CREs with strong DSX binding sites that regulate *dsx* expression in the developing wing.

In addition to the spike of widespread *dsx^H^* expression in early pupal wings, DSX^H^ becomes uniquely expressed in regions of the wing that will become white in mid-pupal wings^15^. Interestingly, only one *H*-specific CRE was even accessible in mid-pupal wings (Supplementary Figure 4). Instead, we found three conserved CREs were DB by DSX between the sexes in both alleles, and 10 additional conserved CREs DB specifically between *HH* males and females in mid-pupal wings (Supplementary Figure 11). We therefore predict that *H*-specific CREs and DSX binding may be required early in development to trigger the developmental switch, but that conserved CREs are differentially used to sustain the mimetic color pattern program throughout development.

## DSX directly regulates a small number of genes in early pupal wings to switch on mimetic color pattern development

Antibody stains showed that DSX^H^ expression never fully pre-figured the adult mimetic pattern, suggesting that DSX^H^ initiates mimetic pattern development in early pupal wings but quickly becomes decoupled from it^15^. We expected that DSX^H^ initiated the mimetic program by directly regulating one or more downstream genes in early pupae that execute the mimetic program^11,12,15^. To identify these direct targets and characterize the consequences of DSX binding on chromatin accessibility and gene expression, we directly compared genome-wide patterns of DSX CUT&RUN, ATAC-seq, and differential gene expression (DE) between mimetic and non-mimetic females (**Fig 4**).

**Fig 4.**
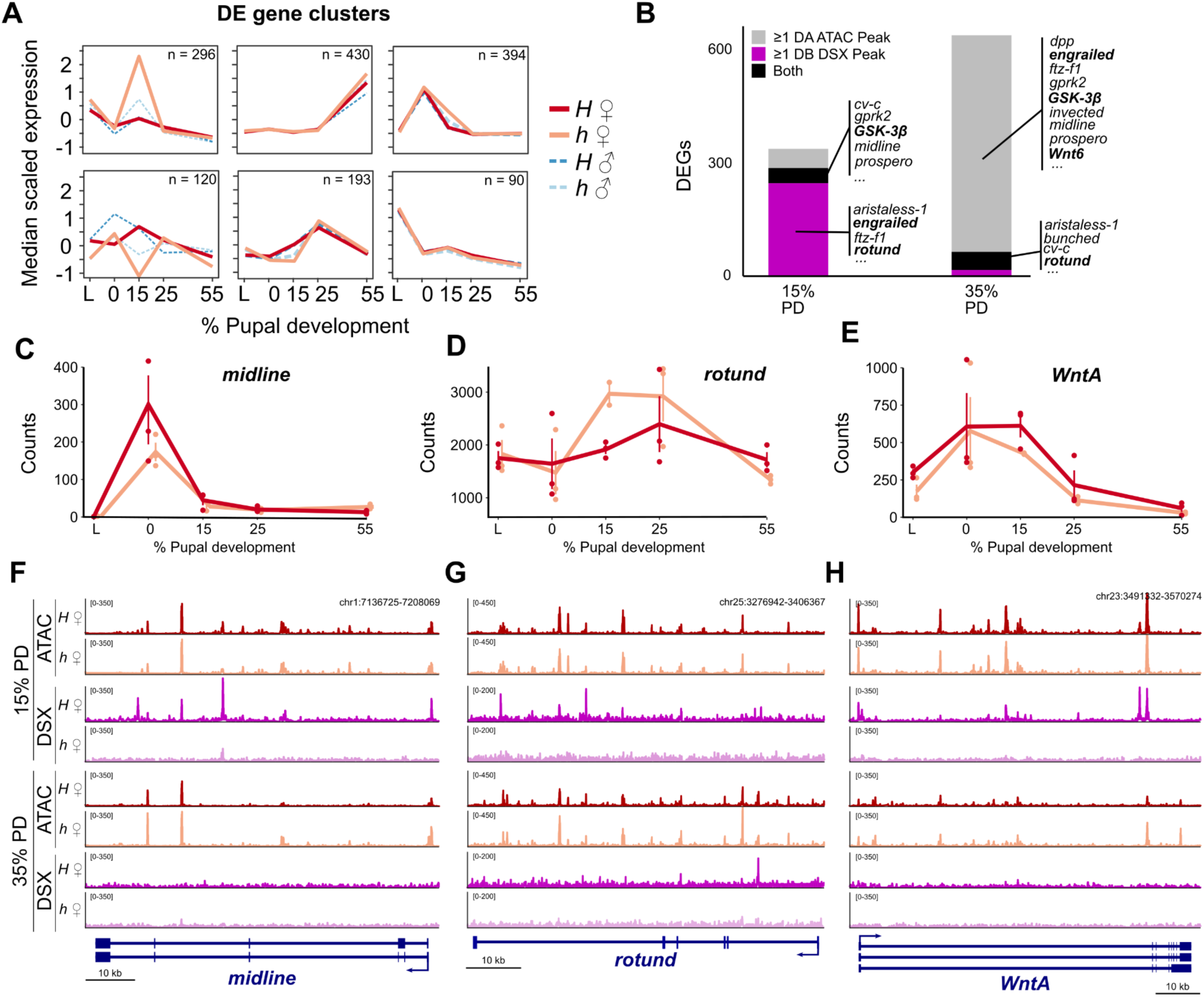
Acute and long-term consequences of DSX binding in early pupal wings. **A,** Median expression profiles of clustered differentially expressed (DE) genes between mimetic and non-mimetic females. Hierarchical clustering was performed using Mclust, using BIC to choose *k*^33^. L: 5th instar larva. **B**, Differential DSX binding and differential ATAC peak accessibility in DE genes. Select genes known to be involved in wing color pattern development, transcription, or signaling are shown. Genes with experimental evidence of their involvement in the mimicry switch are bolded ^15,34^. **C-E,** Expression profiles of direct targets of DSX regulation in mimetic and non-mimetic females. **F-H,** ATAC and DSX CUT&RUN profiles in early and mid pupal development for the genes shown in **C-E**. Gene models are shown with blue boxes (exons) and lines (introns) below. Full dataset can be found in Supplementary Table 13.

We first identified DE genes by re-analyzing wing RNA-seq data from five developmental stages^15^, then intersected those genes with our DSX and ATAC peak sets. Consistent with our previous work, 1523 genes were DE between mimetic and non-mimetic females across development, with the majority (54.8%) of those genes being DE specifically in early pupal wings (**Fig 4A**; Supplementary Figure 12; Supplementary Table 13). In parallel, we found that 3640 (35.3%) DSX peaks in 2187 genes were DB between early pupal mimetic and non-mimetic females, including 248 peaks in 148 early pupal DE genes (**Fig 4B**). These potentially direct targets include *rotund* (a TF previously implicated in the mimicry switch in *P. polytes*^34^), the T-box TF *midline*, the homeobox TF *aristaless-1*, and at least eight other DNA-binding proteins (**Fig 4B**; Supplementary Table 13). The small number of genes that DSX appears to directly regulate in *P. alphenor* wings is consistent with work in *Drosophila* that showed DSX binds many targets genome-wide, but affects expression of a small number of those genes, presumably due to the availability of appropriate co-factors^35^. By mid-pupal development, only 557 (5.4%) DSX peaks were DB, with just 46 peaks in 41 DE genes, including *rn* and *al-1*.

Patterns of chromatin accessibility were opposite those of DSX binding. While 1.7% of ATAC peaks were DA between early pupal female wings, 27.7% were DA in mid-pupal female wings, suggesting that DSX binding early in development alters the wing regulatory landscape throughout development (Supplementary Figure 13). More specifically, 5.6% of DE genes contained DA peaks in early pupal wings, yet 40.8% contained DA ATAC peaks in mid-pupal wings, including genes known to be involved in the *dsx* mimicry switch and to specify color patterns in other butterflies, including *invected, engrailed,* and *al-1* (**Fig 4**)^15^. Thus, while DSX appears to directly regulate few genes early in pupal development, its effects are propagated to later stages while being decoupled from DSX binding itself.

## Discussion

Our results provide several new insights into the structures and evolution of supergenes that have remained hidden by the extensive LD and complexity that characterize many of the best-characterized supergenes. By experimentally identifying multiple functional elements that are required for the *dsx* supergene’s function, we provide support for predictions that inversions and other mechanisms of recombination suppression link distant functional loci together^16^.

Second, our results suggest that the *dsx* supergene originated via the gain of a novel autoregulatory element(s) that enabled positive reinforcement of *dsx^H^* expression in the early pupal wing, causing a spike in *dsx* expression that initiates the mimetic pattern program^12,15^. We predict that subsequent gain of additional autoregulatory elements helped to refine the novel allele’s expression pattern, and that selection for mimicry favored the evolution of an inversion that linked these novel autoregulatory elements together.

Importantly, autoregulation could provide a simple route to supergene evolution. Although we have discussed autoregulation in its most direct and narrow sense - one gene regulating itself - there are many pathways by which genes could regulate themselves^36^. Auto-regulation could play an identical role in the evolution of multi-gene supergenes, where key TFs regulate their own and/or nearby gene expression. Recombination suppression and subsequent divergence between alleles, in CREs or protein-coding sequences, would then refine the supergene alleles’ functions. Beyond supergenes, our findings provide novel insight into the genetic mechanisms by which pleiotropic genes are co-opted into new developmental roles.

Auto-regulation may avoid many of the deleterious pleiotropic effects from ectopic expression because it can only occur in tissues or stages where the gene is already expressed. Auto-regulation, particularly positive reinforcement, may increase the probability that novel alleles are dominant and therefore immediately exposed to natural selection.

Finally, our results suggest that epistatic modifier loci may evolve by gaining direct regulation by the supergene. All *P. polytes* develop tails on their hindwings, but only mimetic female *P. alphenor* develop tails. Thus, a modifier locus has evolved specifically in *P. alphenor* that responds to the mimetic allele to control the presence of tails^21^. Our results suggest that such a modifier could evolve via the gain of direct regulation by DSX. High DSX expression in the mimetic wing can turn on the tail modifier, while low DSX expression in the non-mimetic wing is insufficient to turn on the tail modifier. Such unlinked modifiers can evolve to improve mimicry because they are inactive, or at least not regulated by DSX, in the non-mimetic wing. The small number of direct DSX targets are therefore prime candidates for characterizing the identifies and molecular functions of these epistatic modifiers, and could provide additional insight into how supergene functions are refined.

## Acknowledgements

We thank the University of Chicago greenhouse staff and the University of Chicago Functional Genomics Facility (RRID:SCR_019196) for research support; the University of Chicago’s Center for Research Informatics for computational support.

## Author contributions

NWV - Conceptualization, Investigation, Visualization, Writing - original draft, Writing - review & editing; SIS - Conceptualization, Investigation; CLF - Investigation; DM - Investigation; WL - Investigation; MRK - Conceptualization, Funding acquisition, Writing - review & editing

## Data Availability

Illumina sequencing data is publicly available in the National Center for Biotechnology Information (NCBI) under BioProject PRJNA1062051. Genome assemblies, annotations, R projects, and full analysis results are publicly available in Dryad accession <u>10.5061/dryad.tx95×6b75</u>.

## Materials and Methods

### Butterfly care

*Papilio alphenor* pupae were purchased from Philippines breeders and allowed to emerge in the University of Chicago greenhouses. New adults were sexed, labeled with a unique number on the forewing, and the sexes separated until use. We determined each individual’s *dsx* genotype using DNA from a single leg and a custom TaqMan (Life Technologies, USA) assay. We set up crosses between multiple virgin adults carrying the desired alleles in 2m^3^ mesh cages, allowed them to mate, and provided *Citrus* (Meyer lemon) shrubs for oviposition. Adults were fed Bird’s Choice artificial nectar and supplied with blooming *Lantana*. Pre-pupae were collected and placed into labeled boxes in an incubator set to 25°C, 16h:8h light/dark cycle, and constant 65% humidity. Pupal development takes approximately 15 days under these conditions; experiments focused on two days and five days after pupation (15% and 35% pupal development, respectively).

### Genome sequencing and assembly

We extracted high molecular weight (HMW) genomic DNA from thorax of freshly killed *P. alphenor* females that were homozygous for the *h* or *H* alleles using the QIAgen GenomicTip G- 100 kit (QIAgen, USA). Extractions followed the manufacturer’s instructions, except we incubated chopped fresh tissue in lysis buffer and proteinase K overnight at 50°C and shaking at 200 rpm before purification. We then constructed Oxford Nanopore sequencing libraries using the ONT Ligation Sequencing Kit (LSK-110) and eliminated reads <10 kb using the PacBio SRE XS kit before sequencing on a MinION Mk1b and R9.4.3 flow cells to 30X - 40X coverage.

We called bases using Guppy and super high quality base calling (dna_r9.4.1_450bps_sup.cfg), then assembled the mimetic *H* and non-mimetic *h* genomes separately using these raw reads and Flye v2.9.1 with default settings with expected genome size set to 250 Mb. The initial Flye assemblies were each polished using the Guppy basecalls and Medaka v1.7.2 (medaka_consensus) with the appropriate error model (r941_min_sup_g507). We then purged duplicates using purge_dups v1.2.5.

We then scaffolded contigs together into 31 chromosomes using Hi-C data (see below) and the 3d-dna pipeline^37^. We identified the Z and W chromosomes by analyzing coverage of re-sequencing data from five males and five females^11^. The remaining 29 chromosomes were ordered and labeled by decreasing size. We assembled the *alphenor* mitochondrial genome using NOVOplasty v4.2^38^ using sequencing data from SRR1108726 and the RefSeq mtDNA assembly for *polytes* (NC_024742.1) as the seed sequence. This resulted in a single circularized sequence of 15,247 bp. We added this sequence as chrM to each assembly.

We identified repeat sequences in the mimetic assembly using RepeatModeler, then used our custom library, the RepBase 20181026 “arthropoda” database, and Dfam 20181026 database to identify and mask repeats genome-wide in both assemblies using RepeatMasker. Finally, we hard masked regions in the nuclear genomes with homology to chrM (blastn e-value < 1e-50).

We sequenced and assembled a new *Papilio nephelus* genome following the same protocol, using DNA from a single random female. *Papilio nephelus* is a sexually monomorphic, non-mimetic species that diverged ∼15 mya from the *P. alphenor* lineage. Final assembly statistics can be found in Supplementary Table 14.

### Annotation

We annotated the mimetic assembly using EvidenceModeler 1.1.1^39^. We first assembled a high-quality transcript database using PASA ^39^, SE50 data generated in VanKuren et al. ^15^, and PE100 and SE50 data from Nallu et al.^40^. After adapter trimming, we performed *de novo* and genome-guided assembly using Trinity v2.10.0^41^ and genome-guided assembly using StringTie v1.3.3^42^. RNA-seq data was also mapped to the mimetic *alphenor* assembly using STAR 2.6.1d^43^, and the resulting alignments used to generate genome-guided assemblies with Trinity and StringTie 1.3.1^42^. We combined *de novo* and genome-guided assemblies using PASA 2.4.1^44^. Evidence for protein-coding regions came from mapping the UniProt/Swiss-Prot (2020_06) database and all Papilionoidea proteins available in NCBI’s GenBank nr protein database (downloaded 6/2020) using exonerate^45^. We identified high-quality multi-exon protein-coding PASA transcripts using TransDecoder (transdecoder.github.io), then used these models to train and run Genemark-ET 4^46^ and GlimmerHMM 3.0.4^47^. We also predicted gene models using Augustus 3.3.2^48^, the supplied *heliconius_melpomene1* parameter set, and hints derived from RNA-seq and protein mapping above. Augustus predictions with >90% of their length covered by hints were considered high-quality models. Transcript, protein, and *ab initio* data were integrated using EVM with the weights in Supplementary Table 15.

Raw EVM models were then updated twice using PASA to add UTRs and identify alternative transcripts. Gene models derived from transposable element proteins were identified using BLASTp and removed from the annotation set. We manually curated the *dsx* region and key color patterning genes. The full annotation comprises 20,674 genes encoding 26,074 protein-coding transcripts, containing 97.2% complete and missing 1.9% of endopterygota single-copy orthologs according to BUSCO v5 and OrthoDB v10. We functionally annotated protein models using eggNOG’s emapper-2.0.1b utility and the v2.0 eggNOG database^49^. We used liftoff ^50^ to transfer these annotations to the non-mimetic reference genome assembly.

Downstream analyses used sequence, transcripts, and proteins from only the 31 main chromosomes and chrM.

### Hi-C sequencing and analysis

We performed Hi-C on developing hindwings using Dovetail Genomics’ (USA) Omni-C kit. Hindwings were dissected from staged pupae then snap frozen in liquid nitrogen and stored at - 80°C until use. Wings from three individuals were pooled and pulverized in liquid nitrogen before proceeding with fixation and proximity ligation according to the manufacturer’s protocol. We performed Hi-C separately on hindwings from males and females homozygous for each *dsx* allele at two and five days after pupation. All eight libraries were pooled and sequenced 2 x 150 bp on an Illumina NovaSeq 6000 S1 flow cell at the University of Chicago Functional Genomics Facility (RRID:SCR_019196). We trimmed adapters and low-quality regions using Trimmomatic 0.39 ^51^, then used Juicer v1.6 ^37^ and bwa 0.7.17 ^52^ to map, sort, and de-duplicate reads.

We used these merged_nodups.txt files as input to the 3d-dna pipeline for genome assembly (above). Separately, we used hicExplorer ^53^ to identify TADs in each sample and to plot results used in Fig 1. Juicer output files were converted to.h5 format using hicConvertFormat and 5 kb resolution (except mfp6 and mmp6, which used 15 kb resolution due to their lower quality), normalized using hicCorrectMatrix and and the Knight-Ruiz method ^54^, then used to call TADs with hicFindTads and default settings except “--correctForMultipleTesting fdr --minDepth 20000 --maxDepth 50000 --step 10000”. Samples mfp6 and mmp6 used settings --minDepth 60000 -- maxDepth 150000 --step 30000 to account for their lower resolution.

### Genome blacklists

We generated genome blacklists by identifying low-mappability regions using genmap v1.3.0 ^55^. We calculated mappability for 50-mers, allowing for 1 mismatch (-k50-e1), then identified low mappability regions as those with mappability < 1. We merged low map regions within 300 bp of each other using bedtools merge, then kept regions > 100 bp. We combined these regions with all regions with mtDNA homology (identified using BLASTn with E < 1e-50) into a single blacklist. This was performed separately for the mimetic and non-mimetic genome assemblies.

### ATAC-seq

We performed ATAC experiments following Lewis and Reed^56^ and Buenrostro et al.^24^ with minor modifications. Wings were dissected in room temperature PBS then immediately transferred into ice cold sucrose buffer in 2 mL dounce homogenizers. Tissues were dissociated using 30 - 50 strokes with the tight pestle, then transferred to cold 1.5 mL tubes. Cells and nuclei were pelleted by centrifugation for 5 min at 1000 x *g* at 4°C, then resuspended in 200 uL lysis buffer (wash buffer: 10 mM Tris-HCl pH7.5, 10 mM NaCl, 3 mM MgCl2 plus 0.2% NP-40) and lysed on ice for 5 min. Nuclei were harvested by centrifugation for 5 min at 1000 x *g* for 5 min and 4°C, then resuspended in 750 uL ice cold wash buffer and counted. Aliquots of 500,000 cells were pelleted for 5 min at 1000 x *g* and 4°C, then resuspended in transposition mix (25 uL TD buffer, 2.5 uL TDE1 enzyme, 22.5 uL water). Transposition was performed 30 min at 37°C shaking at 1000 rpm, then cleaned up using the Zymo DNA Clean and Concentrator 5 kit. Libraries were amplified 10 cycles before double-sided cleanup (0.5X - 1.8X) with SPRI Select Beads (Beckman-Coulter, USA). We sequenced libraries PE50 on a single lane of a NovaSeq X 10B flowcell, for an average of 30M read pairs per sample.

### ATAC-seq analysis

Raw ATAC-seq sequencing reads were trimmed using Trimmomatic 0.39^51^, then mapped to each reference genome using bwa 0.7.17^52^. Duplicate reads were removed using picard^57^. We assessed sample quality using ATACseqQC 1.22.0 ^58^, and used the cleaned BAM files for downstream analysis. Bigwig tracks for visualization were created and normalized using the RPKM method and deepTools2^59^. We called peaks using F-Seq2 following the authors’ recommendations for ATAC-seq data (-pe-l 600-f 0-t 4.0-nfr_upper_limit 150- pe_fragment_size_range auto) in each sample, then merged peaks from biological replicates using bedtools intersect^60^, requiring reciprocal 20% overlap between at least two replicates.

Finally, we combined sample peaksets using bedtools merge to generate a comprehensive peakset that we used to identify differentially accessible peaks (DAPs) using DiffBind 3.8.4 ^61,62^. We performed all relevant pairwise comparisons between sexes, genotypes, and stages, then corrected *p*-values using the Benjamini-Hochberg method^63^. Only DAPs with global FDR < 0.05 were used in downstream analyses.

We annotated the full ATAC peaksets using annotatePeaks.pl from HOMER 4.11.0 ^31^.

### Peak and peak sequence orthology

We assigned orthology between *h* and *H* ATAC peak sequences and outgroup genomes using BLAST 2.2.24. First, we identified the genome region bounded by *nach* and *UXT* in the *P. polytes* (GCF_000836215.1)^12^*, P. protenor* (GCA_029286645.1)*, P. nephelus,* and *P. bianor* (available at http://gigadb.org/dataset/100653)^64^ genomes by mapping the non-mimetic *P*.

*alphenor nach, dsx,* and *UXT* transcripts to each genome using minimap2 with the “-xsplice” option^65^. We then converted the minimap2 results to GTF format using paftools.js and UCSCtools bedToGenePred and genePredToGtf tools. We extracted and oriented each region, then used blastn to identify *h* and *H* sequences in each target region. We used the following command, for example:

blastn-task blastn-short-evalue 1e-3 \-outfmt “6 qseqid qlen qstart qend sseqid sstart send length \ pident evalue”-max_hsps 1-query papAlpH.peaks.fa-db papBia.fa

We collated BLAST results and *P. alphenor* peak sequences in R, then plotted the results using the gggenomes package^66^. An R project containing the analysis and plotting pipelines can be found in Dryad (AAAAAAAAA).

### CUT&RUN

We assessed genome-wide Dsx binding in homozygous males and females at 15% and 35% pupal development using CUT&RUN^67^, following the protocol described by Meers et al.^32^ with minor modifications. We dissected hindwings in room temperature (RT) PBS, then enzymatically dissociated cells following Prakash and Monteiro^68^. That is, wings were immediately transferred to 750 uL TrypLE Select Enzyme (Gibco, USA) diluted to 5X in PBS, then broken up by gently pipetting 20X with a p1000 pipet set to 500 uL. Tissues were allowed to dissociate at 32°C / 1200 rpm in a thermomixer for 20 minutes, pipetting 5X every 5 minutes. Cells were harvested by centrifugation at 600 x *g* for 3 min, washed twice with 1 mL RT wash buffer (WB), then resuspended in 1 mL RT WB. Cells were bound to concanavalin A-bound magnetic beads (Bangs Labs) for 10 min rotating at RT. Beads were separated using a magnetic stand, then immediately resuspended in ice cold antibody buffer (WB + 0.025% digitonin). Permeabilized cells (500 000 per reaction) were aliquoted to 0.2 mL tubes before adding 0.5 ug primary antibody and nutating overnight at 4°C. Washing, digestion, and purification followed Meers et al. (2019).

We used rabbit anti-Dsx^15^, mouse anti-H3K4me3 (Abcam ab8580), or goat anti-rabbit IgG (Cell Signaling Technologies 45262).

We constructed sequencing libraries using the NEBNext Ultra II DNA Library Prep kit following the protocol outlined in Liu (2021) “Library Prep for CUT&RUN with NEBNext® Ultra™ II DNA Library Prep Kit for Illumina® (E7645) V.2 (https://www.protocols.io/view/library-prep-for-cut-amp-run-with-nebnext-ultra-ii-kxygxm7pkl8j/v2). The key differences between this protocol and the manufacturer’s protocol are lower annealing temperatures during end repair and PCR. Libraries were pooled and sequenced PE50 on a NovaSeq 6000 in a single SP flow cell to yield ∼10M read pairs per sample (Supplementary Table 10).

### CUT&RUN analysis

Raw CUT&RUN sequencing reads were trimmed using Trimmomatic 0.39^51^, then mapped to each reference genome using bwa 0.7.17^52^. Duplicate reads were removed using picard^57^.

Bigwig files for visualization were generated from these filtered BAM files using RPKM normalization in deepTools2^59^. We called peaks using MACS3^69^ for each sample (using IgG tracks as control tracks) and merged peak calls from biological replicates using bedtools^60^, requiring reciprocal 20% overlap between at least two of three replicates. Finally, we combined peaksets from different samples using bedtools merge to generate a comprehensive peakset that we used to identify differentially bound peaks (DBPs) using DiffBind 3.8.4^61,62^. We performed all pairwise comparisons between sexes, genotypes, and stages, then corrected *p*- values using the Benjamini-Hochberg method^63^. Only DBPs with global FDR < 0.05 were used in downstream analyses.

We identified enriched motifs and annotated the full Dsx peakset relative to gene models using HOMER 4.11.0^31^. We also identified transcription factor binding site motifs enriched in *dsx* peaks (summits +- 100 bp) using HOMER’s findMotifsGenome.pl script with default parameters.

To identify potential TFBSs in *Drosophila melanogaster doublesex*, we downloaded ChIP-seq peak calls for all 546 DNA binding proteins assayed by the modENCODE project in whole flies^30^, then intersected those data with the r6.55 *doublesex* gene model (plus 1 kb) using bedtools intersect^60^.

### CRISPR/Cas9

We designed pairs of guide RNAs (gRNAs) to flank target ATAC peaks using Integrated DNA Technology’s (IDT’s) gRNA Design Tool, then purchased as 2 nmol single gRNAs from IDT (Supplementary Table 8). sgRNAs were resuspended to 1 ug/uL (32 nM) in water. Injection mixes with two sgRNAs consisted of 125 ng/uL each sgRNA and 500 ng/uL IDT SpCas9 V3 in PBS. Injection mixes with 3 - 4 sgRNAs consisted of 75 ng/uL each sgRNA and 500 ng/uL Cas9. Reagents were mixed in a PCR tube, incubated at 37°C for 10 min to complex, then stored at-80°C in 5 uL aliquots until use.

We allowed females to lay on a fresh *Citrus* shrub for 1 - 3 hours, then collected eggs for injections. Eggs were aligned on a small strip of double-sided tape on a glass slide. We injected a small amount of injection mix into the bottom of each egg using pulled borosilicate needles.

The double sided tape was then placed directly onto fresh *Citrus* leaves, where the eggs were allowed to hatch and develop. Results are shown in Supplementary Table 9.

### Mosaic G0 genotyping

HMW gDNA was extracted from thorax of mosaic individuals that appeared to be largely mutant using the QIAgen G-20 tip using the protocol described for genome sequencing. Oxford Nanopore sequencing libraries were then constructed using the Ligation Sequencing Kit LSK- 114, and then sequenced 72 hours on a single R10.4.1 flow cell. Raw reads were converted to base calls using GUPPY, then aligned to the mimetic reference genome using minimap2. Deletions were identified manually in IGV.

### RNA-seq analysis

We re-analyzed SE50 RNA-seq data from VanKuren et al. (2023) following their pipeline. Sequencing data are available in NCBI BioProject PRJNA882073. We quantified transcript expression levels using the raw reads and Salmon 1.9.0^70^. We used k-mer size of 23 and quantified against the transcripts from the mimetic *P. alphenor* assembly, using the whole genome sequence as a decoy. We allowed Salmon to correct for GC, positional, and sequence bias. We then loaded gene-level quantifications using tximport^71^ and identified differentially expressed genes (DEGs) between mimetic and non-mimetic females at each developmental stage. We also identified genes with significantly different developmental expression profiles using maSigpro^72,73^ following our previously described protocol^15^, keeping significant genes with fit correlations >= 0.9.

