## Supplementary Figures for "Supergene evolution via gain of autoregulation"

### Supergene evolution via gain of auto-regulation

**Supplementary Text**
**Supplementary Figures 1 - 13**
**Supplementary Tables 1 - 15**

#### Supplementary Text

##### ***Identification of putative causal mutations in CRE CRISPR/Cas9 experiments***

*Attempts to isolate individual alleles via crossing*

We took two approaches to try to identify causative mutations in mutant G0 butterflies. First, we attempted to isolate individual mutant alleles by crossing mosaic G0 mutant butterflies together to produce F1s, following Mazo-Vargas et al. (2022)(*1*). We attempted to isolate mutations from injection mixes S (targeting mim_peak.22692), R (mim_peak.22673), and P (mim_peak.22667). For each injection mix, we crossed multiple mutant (heterozygous *H/h*) females and multiple *h/h* males together, allowed them to mate and lay eggs. We then screened F1 adults for mutant color patterns and genotyped all individuals for their *dsx* alleles. We expected that heterozygous F1 females carrying mutations that disrupted $dsx^H$ function should then develop non-mimetic color patterns, and we would sequence those individuals to identify the exact causative mutations. We never recovered heterozygous females with non-mimetic color patterns despite screening over 300 butterflies from each cross (including 123, 78, and 115 heterozygous females, respectively).

There are at least two possible explanations for these results. First, the mosaic G0 females that we included in these crosses did not carry germline mutations. This is possible given that each female appeared to be “only” 15% - 25% mosaic. Second, disrupting these CREs may have affected *dsx* expression that is required for gametogenesis or development. Unlike color patterning genes like *WntA* or *Optix*, *dsx* is required for proper sexual differentiation and gametogenesis. Whatever the reason, we were unable to isolate individual mutant alleles by crossing.

*PCR-based screening*

As discussed by Mazo-Vargas et al. (2022) and highlighted in a number of additional studies, CRISPR/Cas9 experiments that use multiple sgRNAs may cause large deletions that span hundreds or even thousands of base pairs beyond the target sites (*2*, *3*). Thus, accurate identification of potential mutant alleles must avoid PCR amplification, which could fail to detect deletions whose breakpoints fall outside of priming sites.

However, these studies, and many others, also highlight the need to identify potential causative mutations to ensure that induced mutations do not simply disrupt nearby coding sequences or nearby, and perhaps essential, functional elements.

*PCR-free whole genome sequencing*

We therefore resorted to performing deep sequencing of a handful of mKO G0 females using amplification-free whole genome long-read sequencing with Oxford Nanopore technologies. We performed deep sequencing (~60X-80X) on mKOs from three different injection mixes and multiplexed, shallow sequencing on an additional four individuals. We extracted genomic DNA from thorax of mosaic G0 butterflies using the QIAgen GenomicTip G-100 kit as detailed above, then constructed sequencing libraries using the SQK-LSK114 sequencing library prep kit following the manufacturer’s instructions with the following modifications: 1) end repair and A-tailing were allowed to proceed for 30 min at 20ºC followed by 30 min at 65ºC instead of 5 min each; 2) all bead elution steps were carried out at 37ºC for 15 min shaking at 700 rpm; 3) ligation was allowed to proceed for 1 hour at room temperature; and 4) final library bead washes were performed with short fragment buffer (SFB). Libraries were then prepped and loaded onto a R10.4.1 flow cell following the manufacturer’s instructions and allowed to sequence for 72 hours. We then called bases using GUPPY and the dna_r10.4.1_e8.2_400bps_sup.cfg error model. We then mapped reads to the *P. alphenor* (mimetic) reference genome using minimap2 v2.26 with the “-x map-ont” option and loaded the resulting BAM file into IGV for visualizing read mapping near the target CRE.

It is also important to note that target CREs were generally far from any coding exons. In fact, CRE
Mim_peak.22692 → 4 kb from TSS
Mim_peak.22673 → 6.5 kb from exon 5 (not a part of the female transcripts)
Mim_peak.22670 → 0.5 kb from exon 5 (not a part of the female transcripts)

Mim_peak.22667 → 4.9 kb from exon 5 (not a part of the female transcripts)
Mim_peak.22663 → 14 kb from exon 6

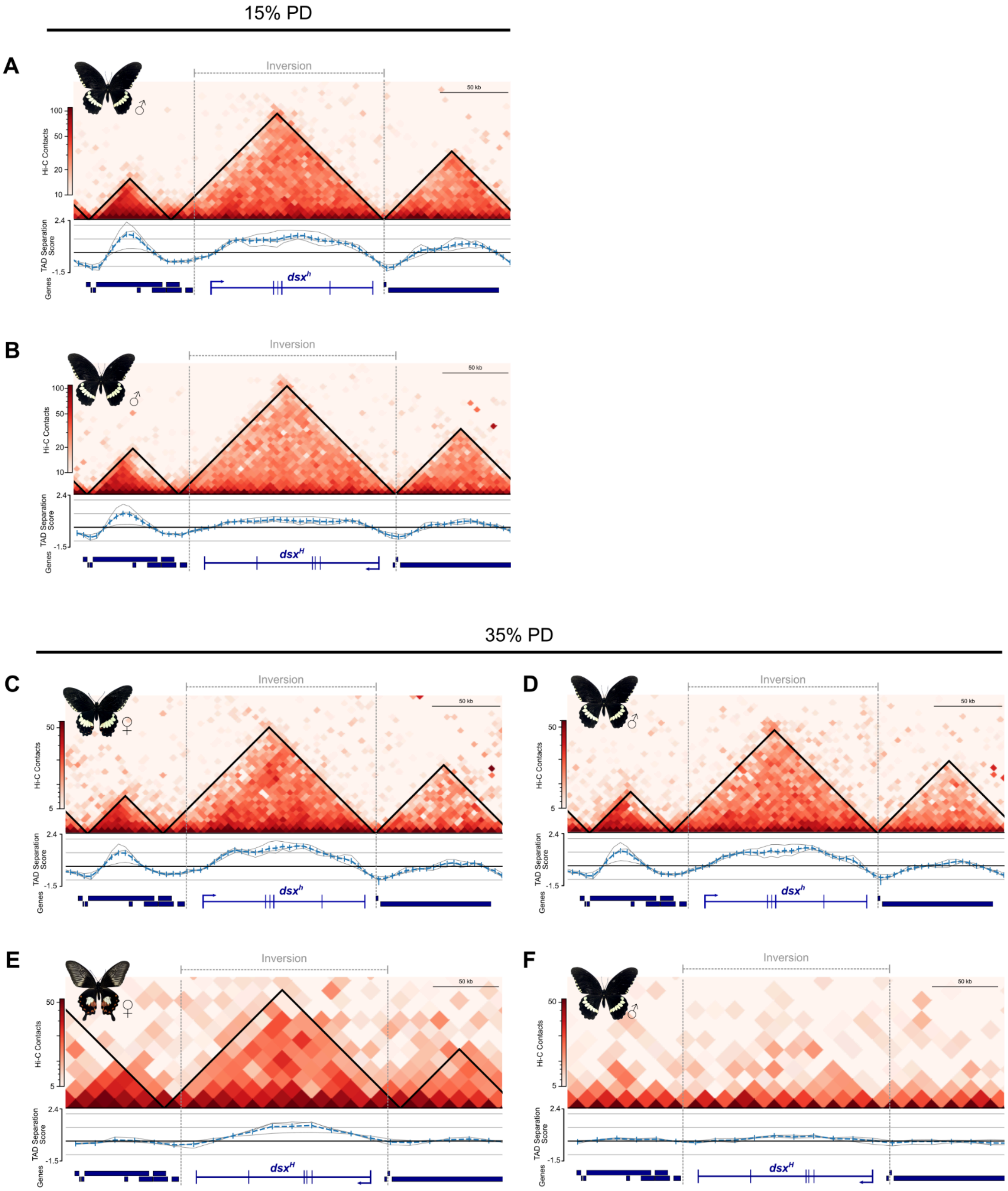


**Supplementary Figure 1. Hi-C contact frequencies and topologically associating domain (TAD) identification in males and mid-pupal female wings.** Supplement to Fig 1C-D. Early pupal (15% pupal development) *h/h* male (**A**) and *H/H* male (**B**) wings. Mid-pupal (35% pupal development) *h/h* female (**C**) *h/h* male (**D**), *H/H* female (**E**), and *H/H* male (**F**) wings. TAD separation scores and genes are shown below each plot, and TADs are outlined by black triangles. Heatmaps in A-D were calculated in 5 kb bins, while E-F were calculated in 10 kb bins due to their lower quality.

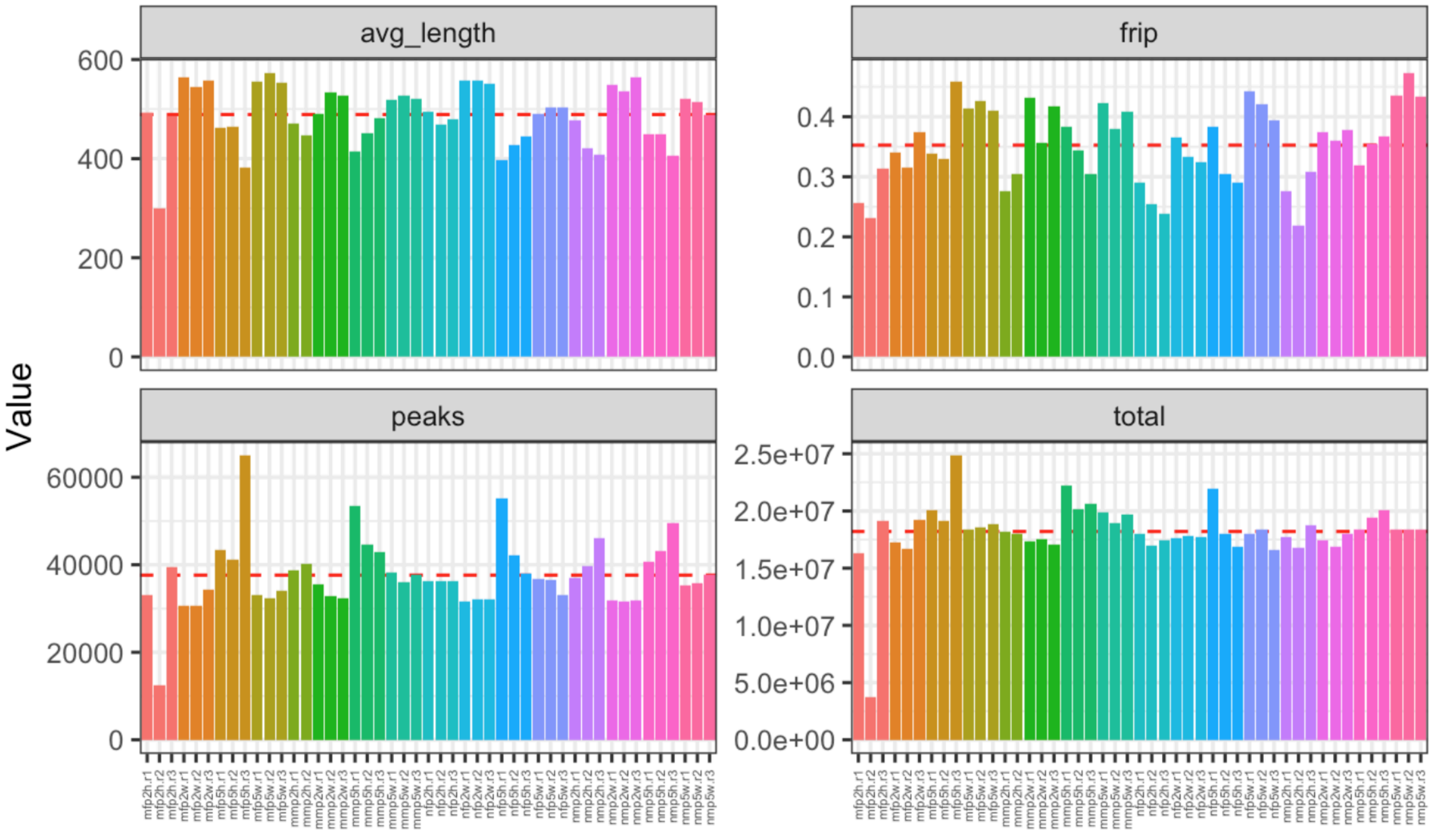


**Supplementary Figure 2. Overview of ATAC data quality.** Clockwise from top left: Average peak length, fraction of reads in sample peaks (frip), total bases in peaks, and total numbers of peaks per sample. Red dashed lines indicate mean values for each metric. Sample information can be found in Supplementary Table 1.

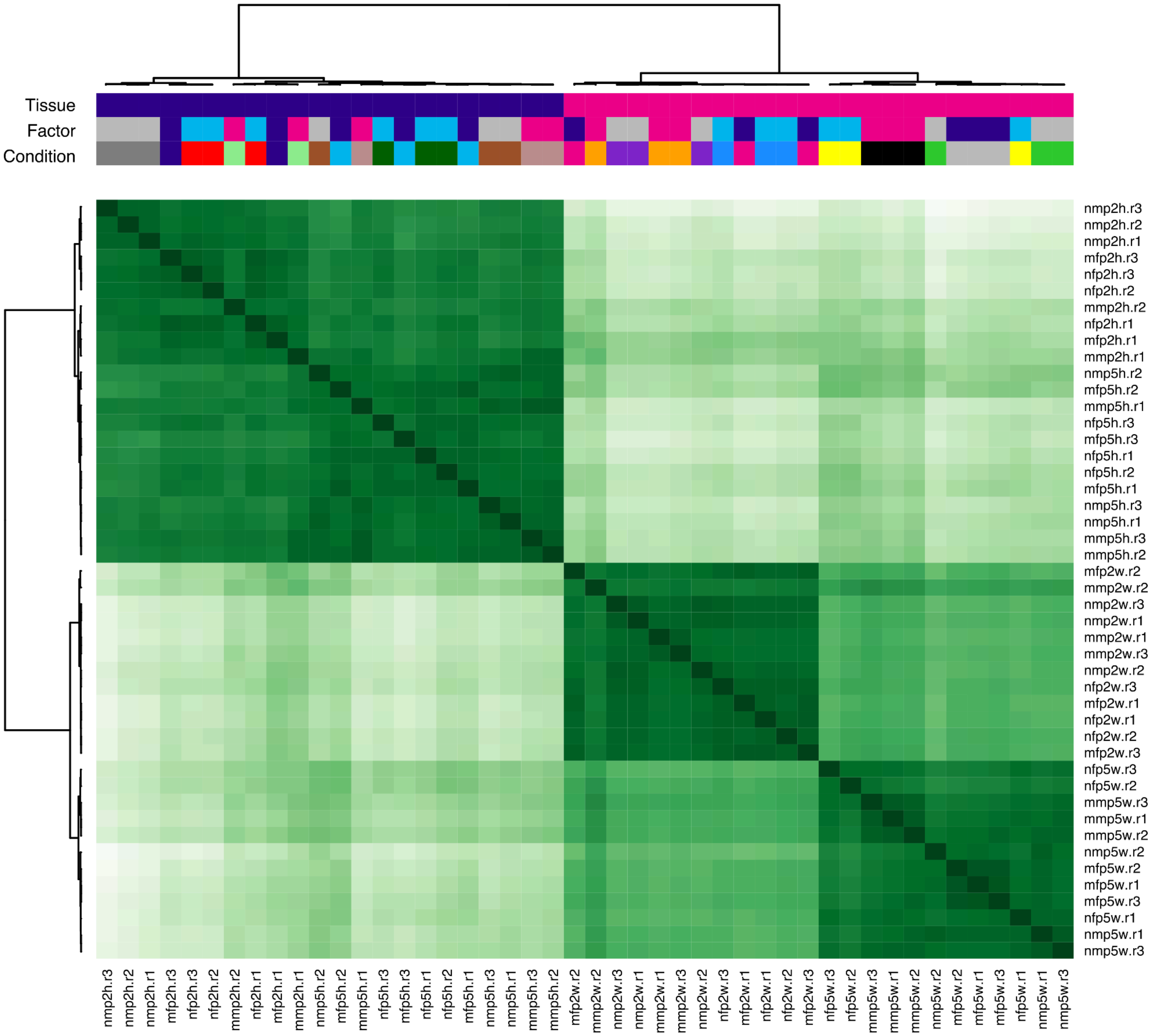


**Supplementary Figure 3. Hierarchical clustering of ATAC samples.** Read depth calculations and plotting were performed using DiffBind. Sample names consist of group-stage-tissue.replicate, where group is *H/H* females (mf) and males (mm) or *h/h* females (nf) and males (nm); stage is p2 or p5 (early or mid-pupal development); tissue is head (h) or wing (w). Primary clustering occurs by tissue, secondary clustering occurs by developmental stage. Sample name and information can be found in Supplementary Table 1.

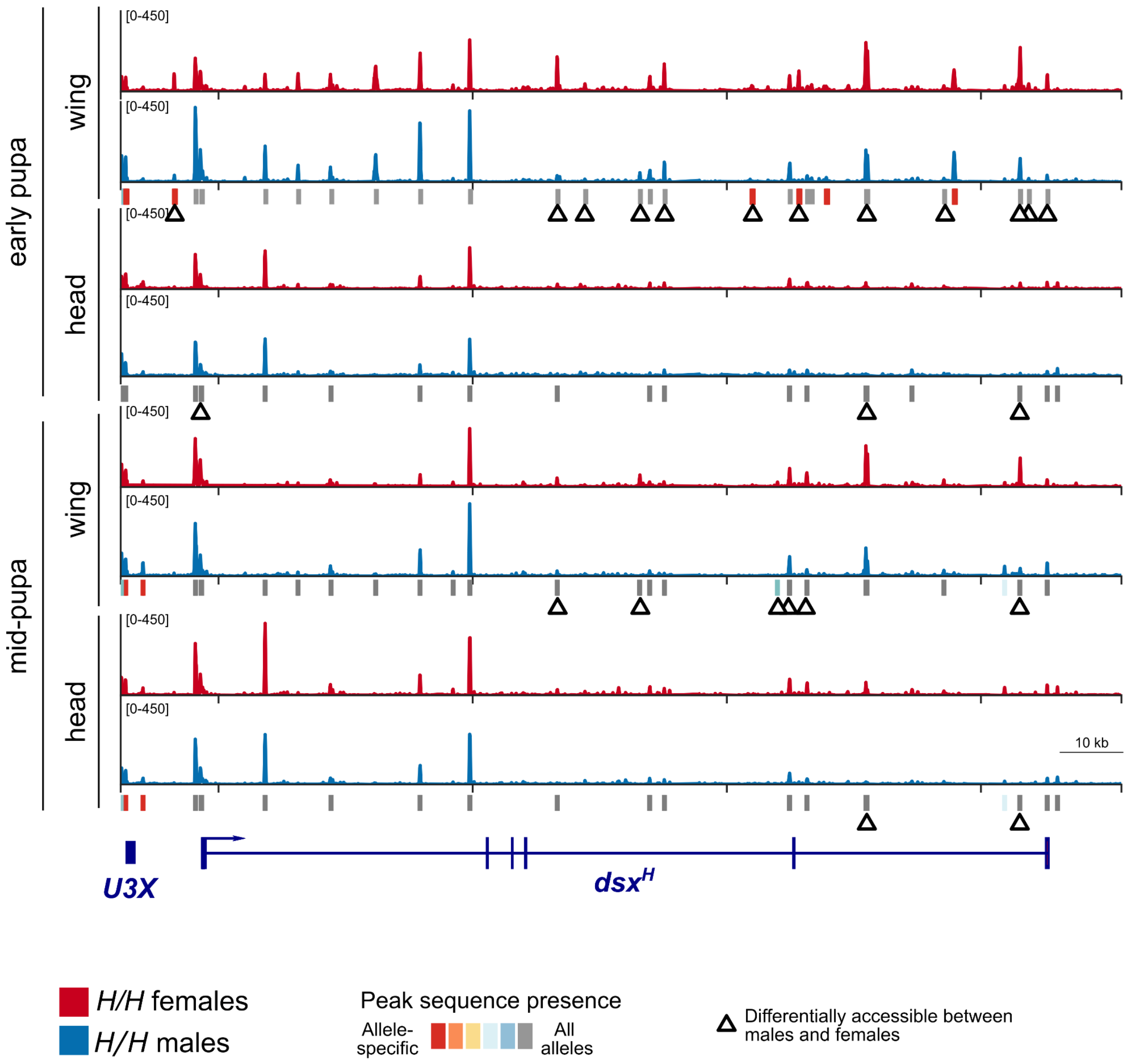


**Supplementary Figure 4. Additional ATAC data in the mimetic *H* supergene allele.** Only the inversion is shown. Tracks for additional replicates and developmental stages, supporting Fig 1D. Peak calls in each tissue/stage are shown below pairs of tracks and colored according to peak sequence presence in other *Papilio* alleles and species. Early pupa: 15% pupal development (p2); mid-pupa: 35% pupal development (p5). Note also that "head" tissues are actually clean brain+retina.

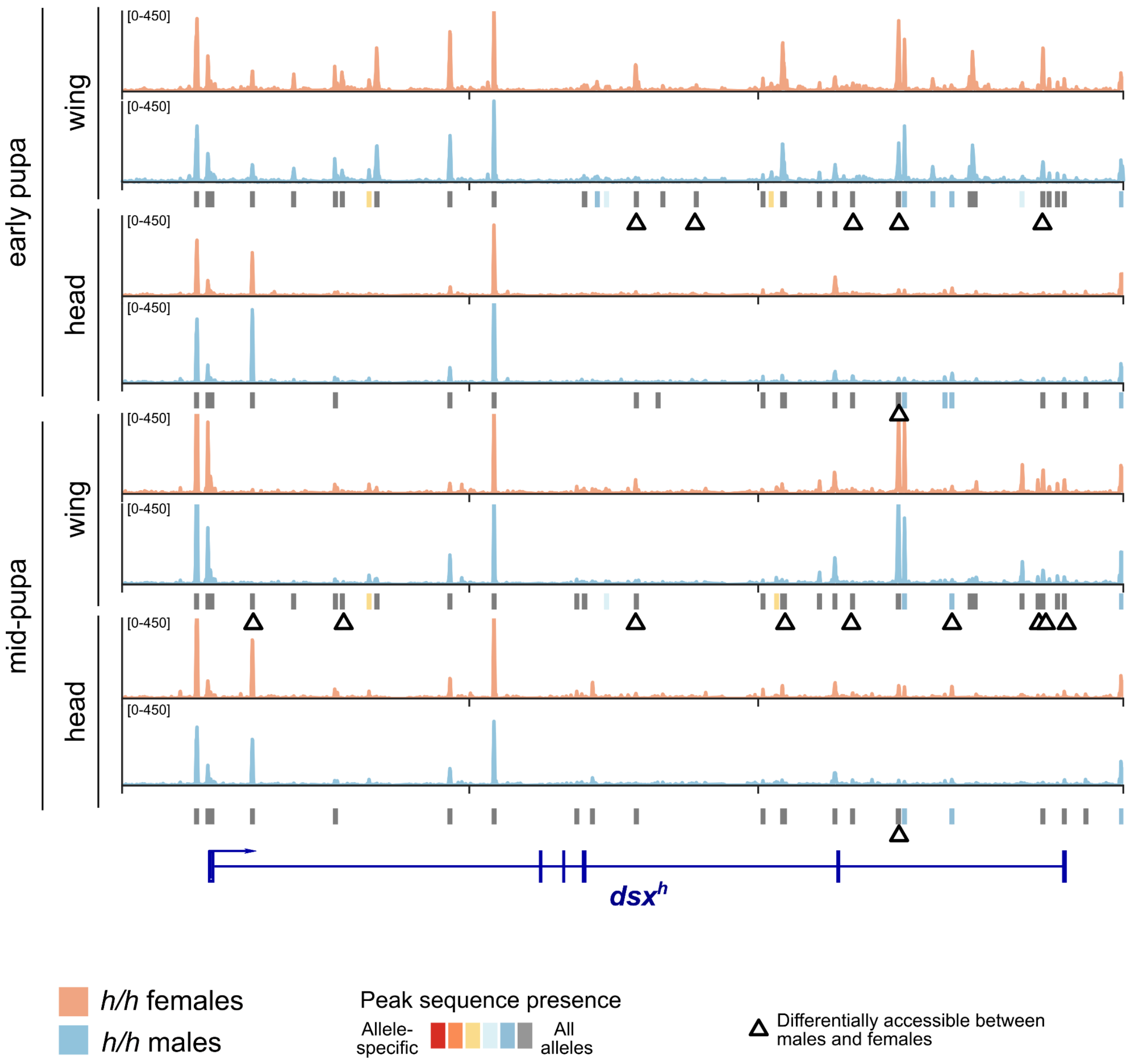


**Supplementary Figure 5. Additional ATAC data in the non-mimetic *h* supergene allele.** Only the inversion is shown. Tracks for additional replicates and developmental stages, supporting Fig 1C. Peak calls in each tissue/stage are shown below pairs of tracks and colored according to peak sequence presence in other *Papilio* alleles and species. Early pupa: 15% pupal development (p2); mid-pupa: 35% pupal development (p5). Note also that "head" tissues are actually clean brain+retina.

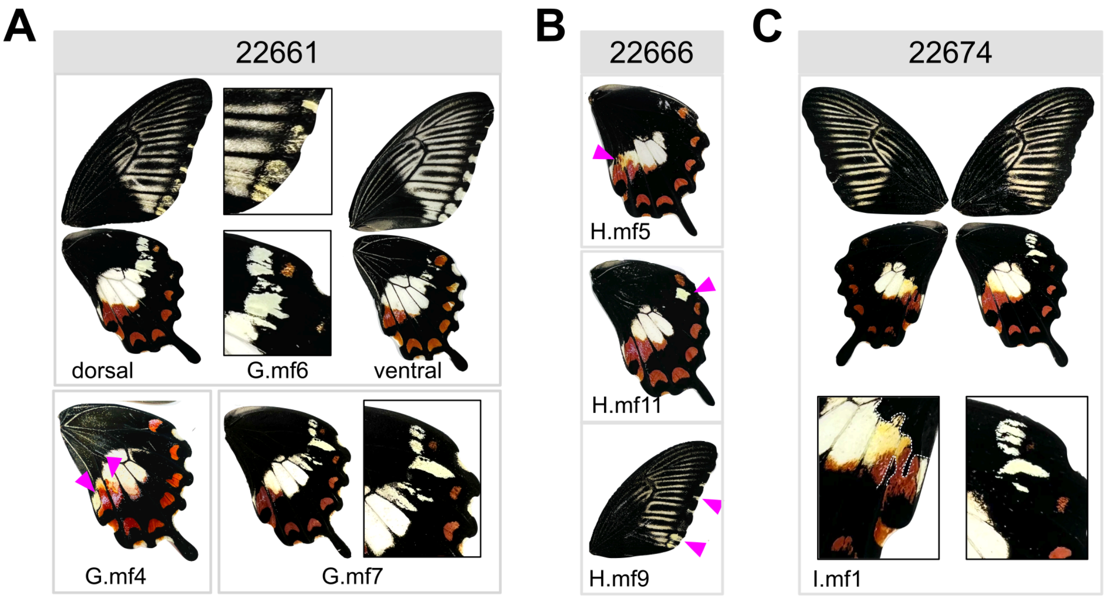


**Supplementary Figure 6. CRISPR/Cas9 KOs of conserved *dsx* CREs.** Supporting information for Fig 2. mKO individuals recovered from injections targeting *dsx* CREs conserved between mimetic and non-mimetic supergene alleles. mKO individuals recovered from injections targeting mim_peak.22661 (**A**), mim_peak.22666 (**B**), and mim_peak.22674 (**C**). Individual identifications (e.g. G.mf6) indicate the injection mix and female recovered. Pink arrows indicate mosaic non-mimetic color patterns. See Supplementary Tables 8 and 9 for sgRNAs and injection mix information.

A

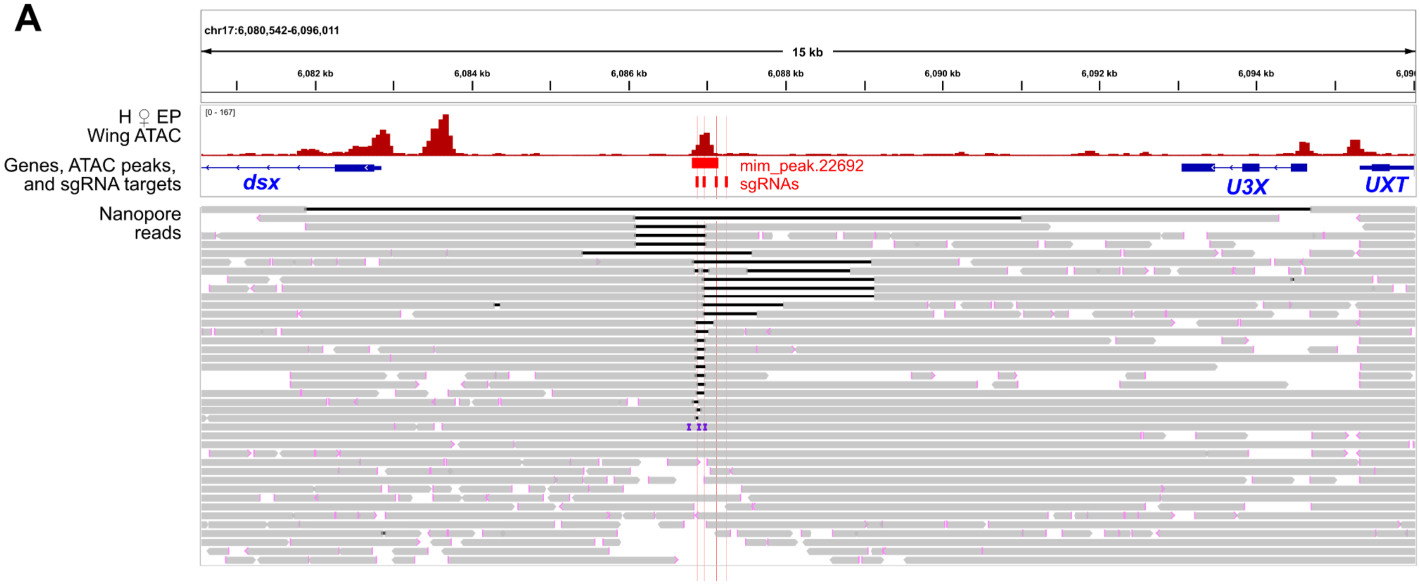


B

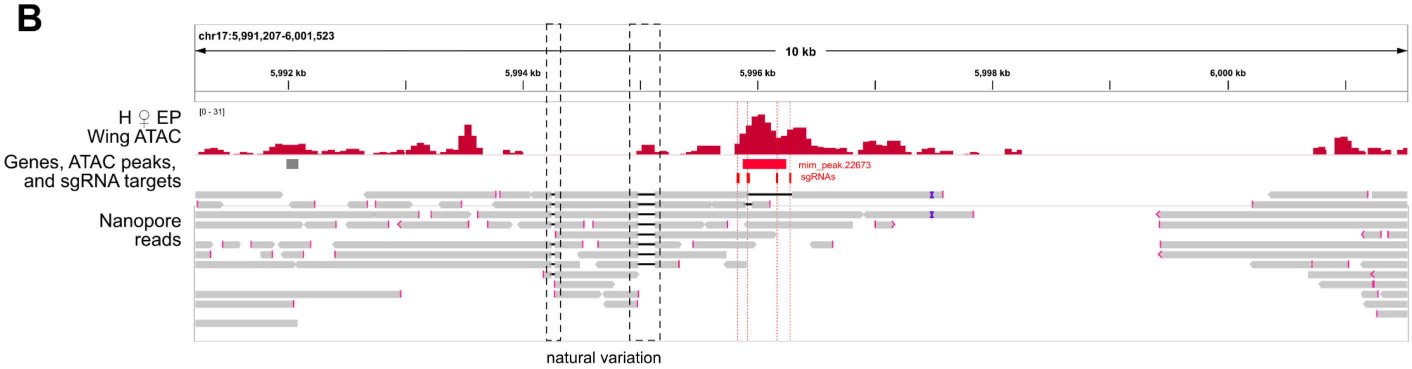


**Supplementary Figure 7. Additional long-read genotyping of CRISPR/Cas9-mediated knockouts of *H*-specific *doublesex* wing CREs.** Each panel shows relevant annotations and all sequencing reads near the target CRE. Reads were aligned to the mimetic reference genome using minimap2, then loaded into IGV to visualize. Small indels and SNPs (<5 bp) are not shown. Red ends of reads indicate soft clipping. Dark lines within a read indicate deletions relative to the reference genome. The ATAC panel shows ATAC-seq data for a single mimetic female early pupal (EP) wing replicate. **A,** Long-read whole genome sequencing of mKO individual S.mf4 (shown in Fig 2C). This CRE resides 4.5 kb upstream of the transcription start site. **B,** Long-read whole genome sequencing of mKO individual R1.mf1 (targeting mim_peak.22673; shown in Fig 2D). This CRE resides in *dsx* intron 4, 6.5 kb upstream of exon 5. These reads also show polymorphic deletions relative to the reference assembly that are nearby but not coincident with gRNA cut sites (dashed boxes).

| Rank | Motif | P-value | log P-pvalue | % of Targets | % of Background | STD(Bg STD) | Best Match/Details |
|---|---|---|---|---|---|---|---|
| 1 * | 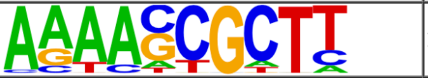 | 1e-11 | -2.672e+01 | 28.95% | 1.36% | 45.5bp (67.9bp) | prd/dmmpmm(Down)/fly(0.663) More Information \| Similar Motifs Found |
| 2 * | 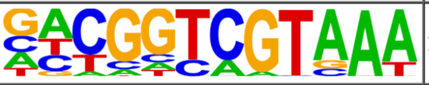 | 1e-11 | -2.592e+01 | 23.68% | 0.70% | 49.6bp (72.1bp) | caudal(Homeobox)/Drosophila-Embryos-ChIP-Chip(modEncode)/Homer(0.786) More Information \| Similar Motifs Found |
| 3 * | 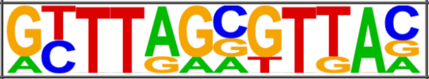 | 1e-10 | -2.526e+01 | 18.42% | 0.27% | 41.3bp (66.7bp) | exd/dmmpmm(Pollard)/fly(0.590) More Information \| Similar Motifs Found |
| 4 * | 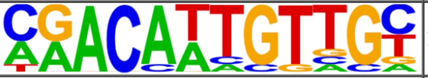 | 1e-9 | -2.261e+01 | 13.16% | 0.08% | 14.2bp (49.6bp) | dsx/dmmpmm(Bergman)/fly(0.756) More Information \| Similar Motifs Found |
| 5 * | 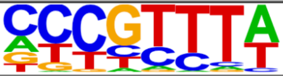 | 1e-7 | -1.830e+01 | 36.84% | 5.75% | 40.9bp (66.7bp) | BH1/dmmpmm(Noyes_hd)/fly(0.690) More Information \| Similar Motifs Found |
| 6 * | 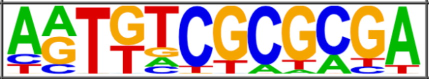 | 1e-6 | -1.561e+01 | 18.42% | 1.09% | 55.1bp (72.8bp) | BEAF-32B/dmmpmm(Pollard)/fly(0.590) More Information \| Similar Motifs Found |
| 7 * | 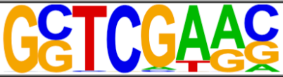 | 1e-6 | -1.451e+01 | 50.00% | 15.15% | 50.1bp (69.7bp) | Deaf1/dmmpmm(Pollard)/fly(0.641) More Information \| Similar Motifs Found |
| 8 * | 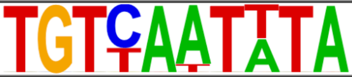 | 1e-4 | -1.114e+01 | 13.16% | 0.82% | 62.6bp (73.2bp) | Vis/dmmpmm(Noyes_hd)/fly(0.736) More Information \| Similar Motifs Found |
| 9 * | 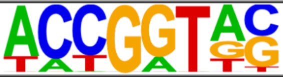 | 1e-4 | -1.079e+01 | 21.05% | 3.16% | 44.3bp (66.4bp) | ovo/dmmpmm(SeSiMCMC)/fly(0.710) More Information \| Similar Motifs Found |
| 10 * | 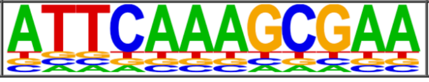 | 1e-2 | -6.424e+00 | 2.63% | 0.00% | 0.0bp (22.1bp) | pan/dmmpmm(Papatsenko)/fly(0.672) More Information \| Similar Motifs Found |

**Supplementary Figure 8. HOMER results from transcription factor binding site enrichment within the mimetic *H* allele.** Target sequences are all *H* allele ATAC peaks, while background sequences were random genomic sequences. Details on HOMER results can be found in the associated help pages (http://homer.ucsd.edu/homer/).

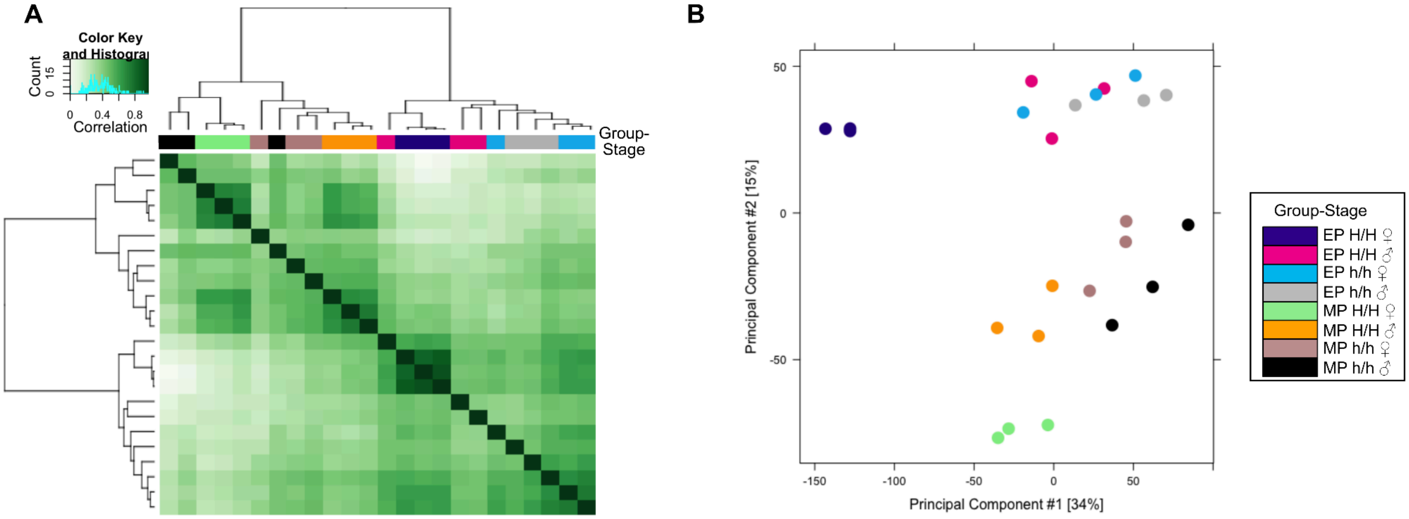


**Supplementary Figure 9. Hierarchical clustering and PCA of DSX CUT&RUN samples.** Read depth calculations, normalization, and plotting were performed using DiffBind. (**A**) Hierarchical clustering of samples based on read depth in all 10,318 DSX peaks. Colors follow the key in **B**, which shows a PC1/2 biplot of sample relationships. EP: early pupa (15%). MP: mid-pupa (35%). Sample name and information can be found in Supplementary Table 10.

| Rank | Motif | P-value | log P-pvalue | % of Targets | % of Background | STD(Bg STD) | Best Match/Details |
|---|---|---|---|---|---|---|---|
| 1 | 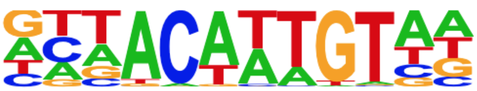 | 1e-744 | -1.715e+03 | 22.60% | 5.48% | 47.8bp (70.6bp) | DMRT6(DM)/Testis-DMRT6-ChIP-Seq(GSE60440)/Homer(0.901) More Information \| Similar Motifs Found |
| 2 | 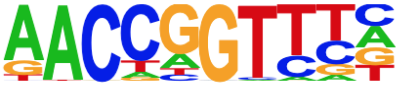 | 1e-300 | -6.924e+02 | 16.13% | 5.85% | 49.8bp (64.2bp) | TFCP2/MA0145.3/Jaspar(0.948) More Information \| Similar Motifs Found |
| 3 | 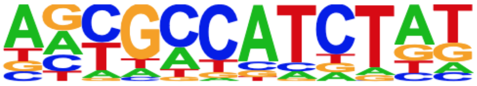 | 1e-289 | -6.665e+02 | 5.34% | 0.69% | 52.2bp (67.0bp) | CTCF/MA0531.1/Jaspar(0.822) More Information \| Similar Motifs Found |
| 4 | 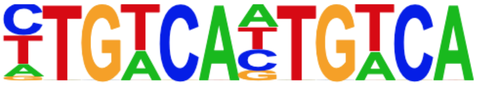 | 1e-263 | -6.065e+02 | 4.48% | 0.51% | 46.2bp (66.8bp) | Tgif1(Homeobox)/mES-Tgif1-ChIP-Seq(GSE55404)/Homer(0.692) More Information \| Similar Motifs Found |
| 5 | 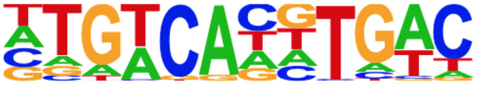 | 1e-189 | -4.360e+02 | 4.98% | 0.98% | 43.6bp (69.5bp) | USF1/MA0093.3/Jaspar(0.846) More Information \| Similar Motifs Found |
| 6 | 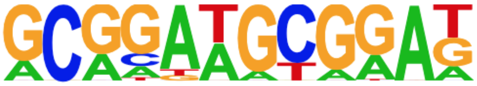 | 1e-132 | -3.059e+02 | 2.10% | 0.22% | 43.1bp (54.0bp) | NRF1(NRF)/MCF7-NRF1-ChIP-Seq(Unpublished)/Homer(0.677) More Information \| Similar Motifs Found |
| 7 | 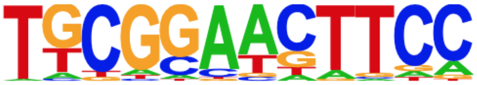 | 1e-98 | -2.273e+02 | 1.31% | 0.10% | 41.2bp (59.6bp) | RDR1/MA0360.1/Jaspar(0.709) More Information \| Similar Motifs Found |
| 8 | 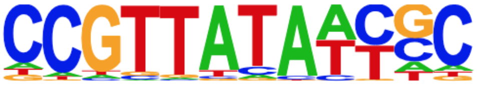 | 1e-97 | -2.253e+02 | 0.75% | 0.02% | 39.7bp (45.5bp) | OVOL2/MA1545.1/Jaspar(0.769) More Information \| Similar Motifs Found |
| 9 | 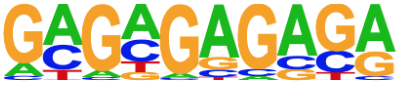 | 1e-91 | -2.106e+02 | 10.63% | 5.51% | 49.4bp (62.7bp) | SeqBias: GA-repeat(0.923) More Information \| Similar Motifs Found |
| 10 | 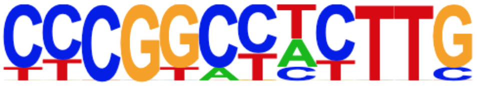 | 1e-86 | -1.999e+02 | 0.95% | 0.05% | 56.5bp (53.7bp) | BHLH112/MA0961.1/Jaspar(0.655) More Information \| Similar Motifs Found |
| 11 | 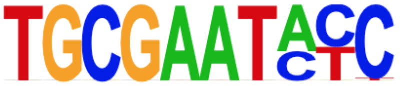 | 1e-68 | -1.580e+02 | 1.03% | 0.10% | 51.1bp (53.6bp) | Ik-1(0.728) More Information \| Similar Motifs Found |
| 12 | 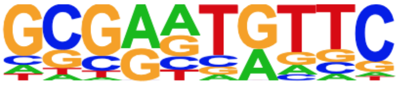 | 1e-48 | -1.124e+02 | 0.92% | 0.12% | 42.4bp (56.8bp) | KAN4/MA1028.1/Jaspar(0.774) More Information \| Similar Motifs Found |

**Supplementary Figure 10. HOMER motif enrichment within genome-wide DSX CUT&RUN peaks.**

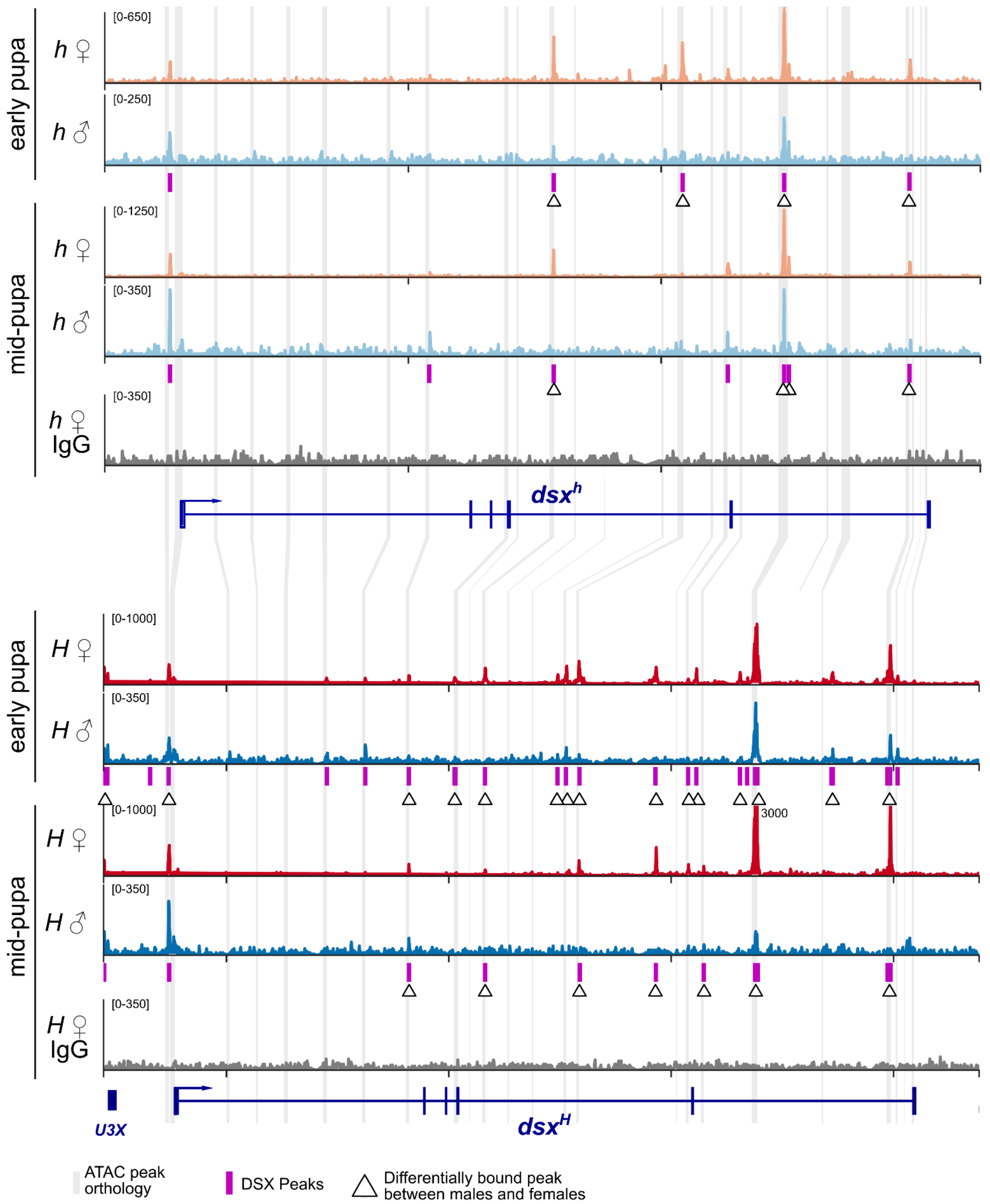


**Supplementary Figure 11. DSX CUT&RUN within the *h* (top) and *H* (bottom; inverted) supergene alleles in pupal wings.** Only the inversion is shown. Normalized CUT&RUN coverage is shown, note differing scales between tracks. The labeled peak in mid-pupal female wings (3000) extends beyond the y axis limits. Sequence orthology is based on ATAC peak orthology shown in Fig 1 and Supplementary Tables 4-5. Differential binding data can be found in Supplementary Tables 11-12.

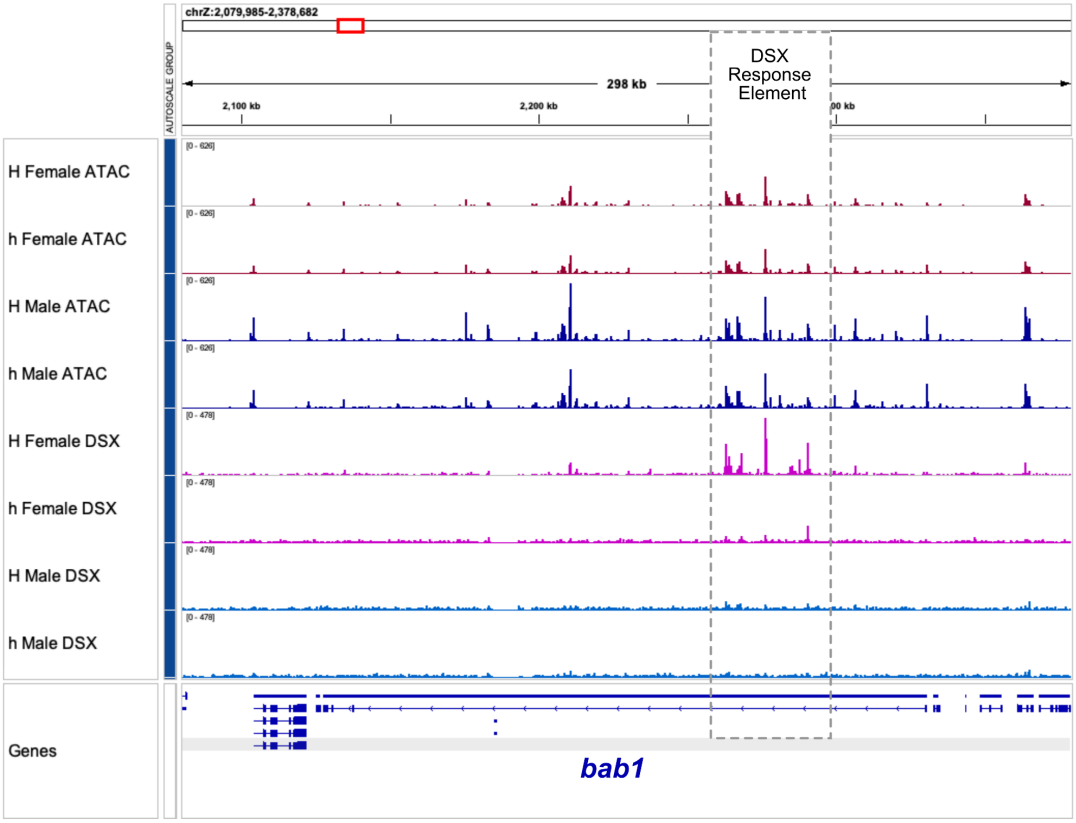


**Supplementary Figure 12. DSX CUT&RUN and ATAC-seq data in *bric-a-brac 1* (*bab1*) in early pupal *Papilio alphenor* wings.** *bab1* is a known DSX target gene in *Drosophila melanogaster,* where DSX binds to the DSX Response Element (*4*, *5*). The orthologous region is boxed here.

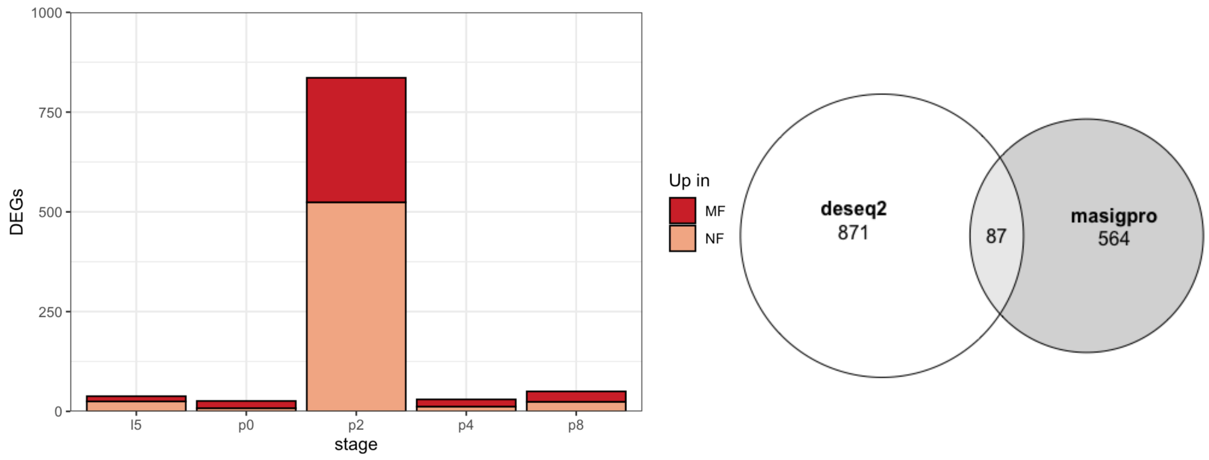


**Supplementary Figure 13. Differential expression between mimetic and non-mimetic females.** DE genes were identified at each stage using DESeq2 or as genes with significantly different expression profiles across development using maSigPro. Left: Stage-specific analysis using DESeq2. Significant genes are those with globally corrected false discovery rate < 0.05. Right: Euler diagram showing the intersection between stage-specific results and maSigPro results. Only significant genes with fit correlations >0.9 were included in the maSigPro set.

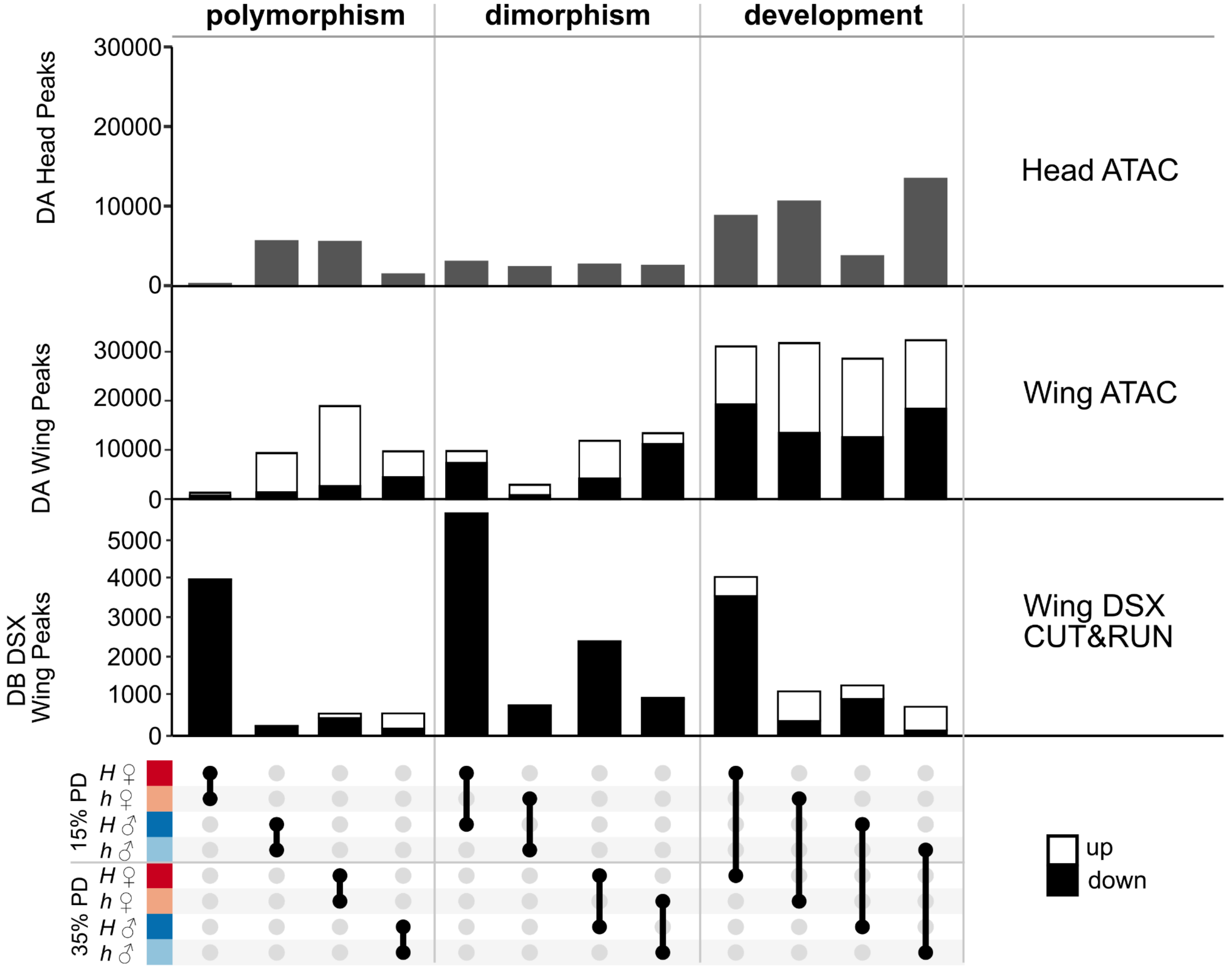


**Supplementary Figure 14. Summary of differential accessibility and differential DSX binding in *Papilio alphenor* pupae for select comparisons.** “Up” is relative to the top point in the ggupset panel (e.g. in the H ♀ - h ♀ comparison, “up” means more reads in H ♀).
